# Adaptation is influenced by the complexity of environmental change during evolution in a dynamic environment

**DOI:** 10.1101/724419

**Authors:** Sébastien Boyer, Lucas Hérissant, Gavin Sherlock

## Abstract

The environmental conditions of microorganisms’ habitats may fluctuate in unpredictable ways, such as changes in temperature, carbon source, pH, and salinity to name a few. Environmental heterogeneity presents a challenge to microorganisms, as they have to adapt not only to be fit under a specific condition, but they must also be robust across many conditions *and be able to deal with the switch between conditions itself*. While experimental evolution has been used to gain insight into the adaptive process, this has largely been in either unvarying or consistently varying conditions. In cases where changing environments have been investigated, relatively little is known about how such environments influence the dynamics of the adaptive process itself, as well as the genetic and phenotypic outcomes. We designed a systematic series of evolution experiments where we used two growth conditions that have differing timescales of adaptation and varied the rate of switching between them. We used lineage tracking to follow adaptation, and whole genome sequenced adaptive clones from each of the experiments. We find that both the switch rate and the order of the conditions influences adaptation, and that switching can both speed up and slow down adaptation, depending on those parameters. We also find different adaptive outcomes, at both the genetic and phenotypic levels, even when populations spent the same amount of total time in the two different conditions, but the order and/or switch rate differed. Thus, in a variable environment adaptation depends not only on the nature of the conditions and phenotypes under selection, but also on the complexity of the manner in which those conditions are combined to result in a given dynamic environment.

## Introduction

How organisms evolve is a fundamental question in biology, and how they adaptively evolve in response to *changing* environments is a question whose answer is central to rational vaccine development^46^, as well as to understanding the evolution of multiple antibiotic resistance^3,22^, the evolution of immune systems^34^, and even heritability^40^. In nature, some environmental changes are predictable, and organisms can evolve responses to such predictable changes. For example, *E. coli* shows asymmetric anticipation of carbon sources, such that in the presence of lactose, *E. coli* anticipates that maltose will soon become available, because this is what has been repeatedly experienced in the mammalian gut. Mechanistically, this is due to lactose modestly inducing the genes required for maltose metabolism^37^. However, when wild *E. coli* are grown under laboratory conditions, which typically lack this selective pressure, this “anticipation” is lost as the strain undergoes domestication^37^. Likewise, circadian clocks are thought to provide a fitness benefit, allowing organisms to adapt physiologically to diurnal changes in light, temperature, and humidity^14^. In Cyanobacteria, the benefit of a circadian clock can only be maintained in the lab by continued exposure to a rhythmic environment^48^. Environmental change may vary based on the frequency of switching, and whether the switching is random or predictable – one way in which organisms can adapt to deal with environmental uncertainty is by bet hedging, whereby by the stochastic switching between different phenotypic states can allow a portion of a population to be more fit under a certain environment^13^. It has been experimentally shown that bet-hedging approaches that resulting in greater average fitness across environments can be engineered^5^ or evolved^1,30^, and there is a rich theory on bet hedging as a strategy to survive in variable environments that switch more rapidly than can be kept up with through mutation and selection alone^8,41^.

Experimental Microbial Evolution (EME^2^; also referred to as Adaptive Laboratory Evolution (ALE)) is a prospective approach to studying adaptive evolution in the laboratory and was first used ~140 years ago^12^. EME has been used to address fundamental evolutionary questions, such as the rate at which beneficial mutations fix^39^, and the influence of both ploidy^39^ and sex^35^ on that rate. High-throughput sequencing has made it possible to establish at high resolution how mutations accumulate in co-evolving lineages, revealing clonal interference, with hundreds or thousands of beneficial lineages competing^20,24–26^, sometimes even with multiple lineages persisting in a quasi-stable state for thousands of generations^16^. While EME has provided many insights into the evolutionary process^(see 11,31 for reviews)^, such experiments have typically been performed in either constant environments (such as the chemostat), consistently fluctuating environments (as in by serial transfer), or in environments where a variable of interest changes monotonically over either time, as in a morbidistat^44^, or over space, as in the megaplate experiment^4^. However, outside of the laboratory organisms are almost never challenged to adaptively evolve in such predictable environments, but rather must cope with variability and stochasticity. To date, only a few EME studies^(see 7 for review)^ have sought to determine either how microbes adapt to unpredictable changes in the environment, or what characteristics of such changes might be important in influencing adaptation. For example, when *Pseudomonas fluorescens* was evolved in variable environments, switching between contrasting carbon sources (xylose and mannose), it was found, contrary to expectation, that populations frequently evolved to be niche specialists, and became adapted to the less favorable carbon source^18^. By contrast, when evolving in a heterogeneous environment containing multiple carbon sources, adaptation converged on the most productive carbon source^19^. In another example, a recent study investigated the fitness of the yeast deletion collection under different time scales of periodic environmental change and showed that some mutants are better at dealing with the environmental switch itself, suggesting that it is possible to evolve genotypes that are adapted to change, *per se*^42^. To date no study has characterized the dynamics of evolution during adaptation to a changing environment or asked specifically how these dynamics might change as a function of the switch rate and strength of selection.

To fill this gap, and to improve our knowledge of how dynamic environments impact the evolutionary process, a systematic (for a given set of environments) exploration of the parameters of dynamic environments is needed, to determine how these parameters affect evolutionary dynamics, and the fitness effects of adaptive mutations across environments. Here we present a series of experiments that explore evolution during switching between two environmental conditions (glucose-containing medium with fluconazole, vs. medium containing ethanol/glycerol with no drug), varying two important parameters: 1) the degree of randomness of the switches between the two conditions, and 2) the consecutive time spent in each condition. Using DNA barcode-based lineage tracking we followed the evolutionary dynamics in 8 different environmental scenarios, investigating the statistics of the evolutionary dynamics, and determining the phenotypic and genotypic characteristics of adaptive mutants arising in each. We found that the speed of adaptation could be either slowed down or sped up depending on the rate of switching between conditions, and that different switching dynamics could lead to the selection of clones with very different behaviors in each environment. Finally, we found that different environmental sequences select for different phenotypic and genotypic outcomes; for example, a randomly switching environment tended to select for generalists, while a consistent strong selection in a non-switching environment selected for specialists.

## Results

### Experimental Design and Overview

We evolved, by serial transfer, barcoded diploid yeast populations in dynamic environments built using two single environment blocks (see Fig. 1 for experimental design), varying two main parameters: i) the time spent in each particular environment, relative to the timescale of adaptation within that environment and ii) the periodicity/randomness of the switching between environments. We defined the timescale of adaptation as the evolutionary time required for a certain fraction of the population to be adaptive within a given environment: for diploid yeast evolving by serial transfer in synthetic complete (SC) medium with 2% glucose + 4 μg/ml Fluconazole (hereafter referred to as “Fluconazole”), ~20% of the population is adaptive after 48 generations, while, in SC medium with 2% glycerol and 2% ethanol (“Gly/Eth”), the timescale of adaptation is much longer: ~15% of the population is adaptive after 144 generations (Fig. 2A and Humphrey, Hérissant et al, in prep.). The timescale of adaptation for these two environments is thus 48 and 144 generations respectively. We designed 8 different evolution experiments (Fig. 1) that combined the Fluconazole and Gly/Eth environments, chosen specifically because of their different timescales of adaptation. The first two experimental sequences were designed so that environmental blocks are periodically switched, with consecutive time spent in each on the order of the time scale of adaptation (periodic_adap1 and periodic_adap2): 144 consecutive generations in Gly/Eth and 48 consecutive generations in Fluconazole. The next two sequences were designed so that blocks were periodically switched at a rate that is 6-fold faster than the previous sequence; thus the consecutive time spent in each environment was 6-times shorter than the time scale of adaptation (periodic_smaller1 and periodic_smaller2): 24 consecutive generations in Gly/Eth and 8 consecutive generations in Fluconazole. We also designed one experiment with random switching between environments, with blocks for which the duration of residence is of the magnitude of the time scale of adaptation (random_adap1), and two experiments that randomly switch between blocks of environment, for which the duration of residence in each environment is less than the time scale of adaptation (random_smaller1 and random_smaller2). We also designed an experiment that combined the two block environments, i.e. SC with 2% glycerol, 2% ethanol and 4μg/ml Fluconazole (Mix), as well as evolved populations in either Gly/Eth, or in Fluconazole, with no switching.

**Figure 1:**
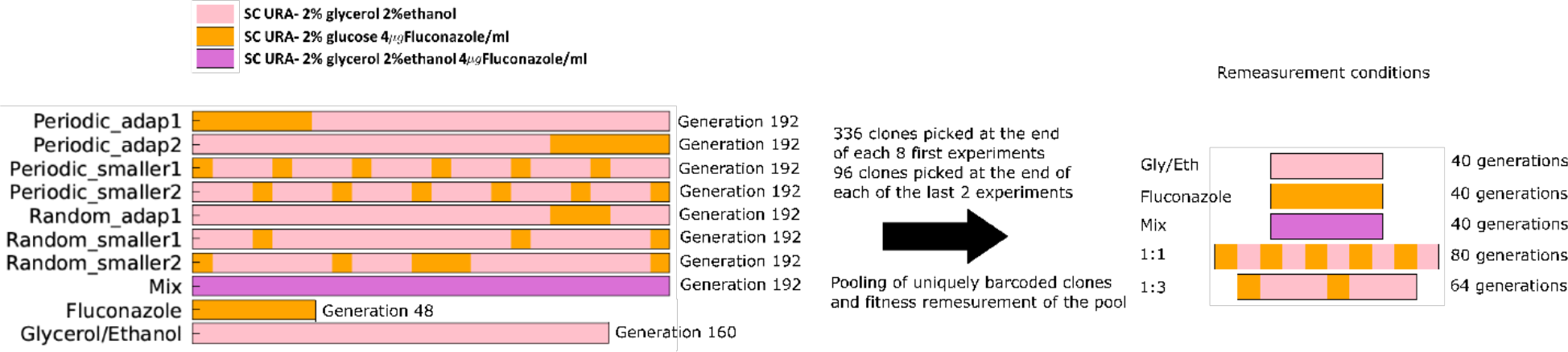
Experimental design. Each experiment with a fluctuating environment was constructed using blocks of 8 generations in Fluconazole and 24 generations in Gly/Eth. At the end of 192 generations, the total time spent in Fluconazole is 48 generations and 144 generation in Gly/Eth, for periodic_adap1, periodic_adap2, periodic_smaller1, periodic_smaller2 and random_smaller2 experiments. By contrast, in random_adap1 and random_smaller1 the total time spent in Fluconazole is 24 generations, with 168 generations in Gly/Eth. Two additional experiments, which did not switch between environments, were also carried out. Clones isolated from each experiment were pooled, and then had their fitness remeasured under 5 different conditions (right).

**Figure 2:**
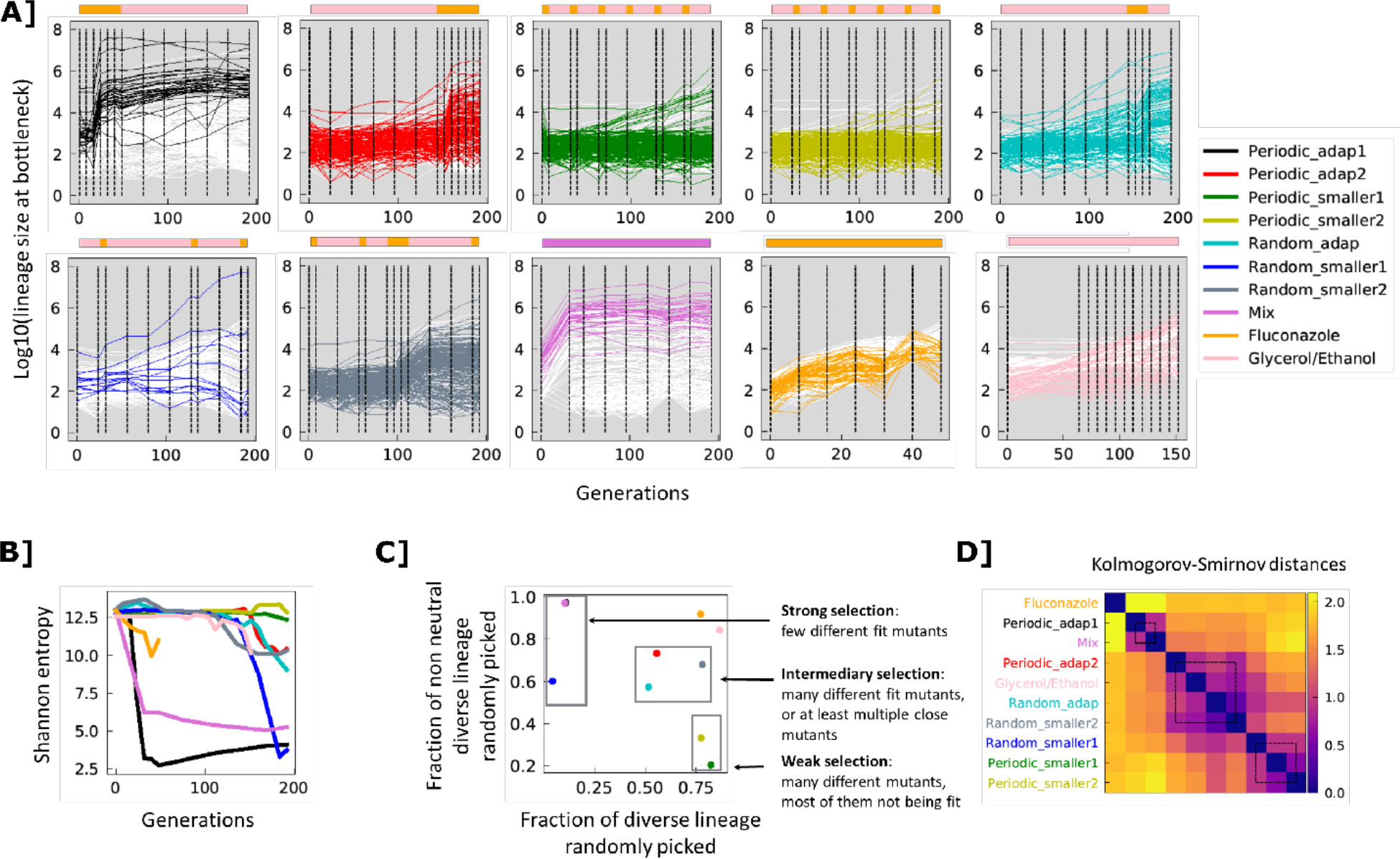
**A] Lineage tracking data for a subsample of 300 lineages for each experiment over the course of 192 generations**. Colored lines correspond to lineages for which single colonies were later isolated; white lines correspond to a randomly selected set lineages which were not sampled for fitness remeasurement. Dashed lines represent sampled time points. Environments are indicated by the color strips above each graph, with colors as in Figure 1. **B] Shannon entropy calculated across all lineages in each experiment. C] Strength of selection**. Barcode diversity and the adaptive fraction for 336 lineages randomly picked from each experiment at generation 192. D] **Similarity of experimental conditions**. Kolmogorov Smirnov distances were calculated between all the experimental conditions based on the fitness remeasurement data. The distance matrix was hierarchically clustered.

Barcoded populations of diploid yeast were then evolved for 192 generations in each of the 8 different sequences of switching environments. The yeast populations contain two barcodes, such that one (BC1, low diversity) encodes the identity of the evolution experiment itself, while the second (BC2, high diversity) is used for lineage tracking within the evolution experiment, to distinguish lineages from one another. We characterized the early stages of adaptation in different dynamic environments using lineage tracking^26^ to follow the population dynamics. We also isolated 336 clones from generation 192 of each evolution, determined their barcodes (see Methods), and pooled unique lineages for which we could recover a barcode sequence. To understand phenotypically how clones from different experiments had adapted to different environmental sequences, we then remeasured the fitness of all lineages in this pool in 5 environments: Fluconazole, Gly/Eth, Mix, 8 generations in Fluconazole and 24 in Gly/Eth (1:3), 8 generations in Fluconazole and 8 in Gly/Eth (1:1) (Fig 3. and SI 1,2). The rationale behind remeasurement in the 1:1 environment was to determine if there has been selection for a phenotype related to their ability to *switch between* environments, instead of fitness in one of the two environment blocks *per se*. Pooled fitness remeasurement experiments were performed in triplicate as previously described^45^, and also included known neutral barcoded lineages, barcoded adaptive yeast from a Fluconazole only evolution and barcoded adaptive yeast from a Gly/Eth only evolution as controls. Neutral lineages in our experiments are defined as lineages that show behavior in the 5 environments similar to known, unevolved neutral lineages (Fig. SI 3). Fitness was determined as described previously^29^.

**Figure 3:**
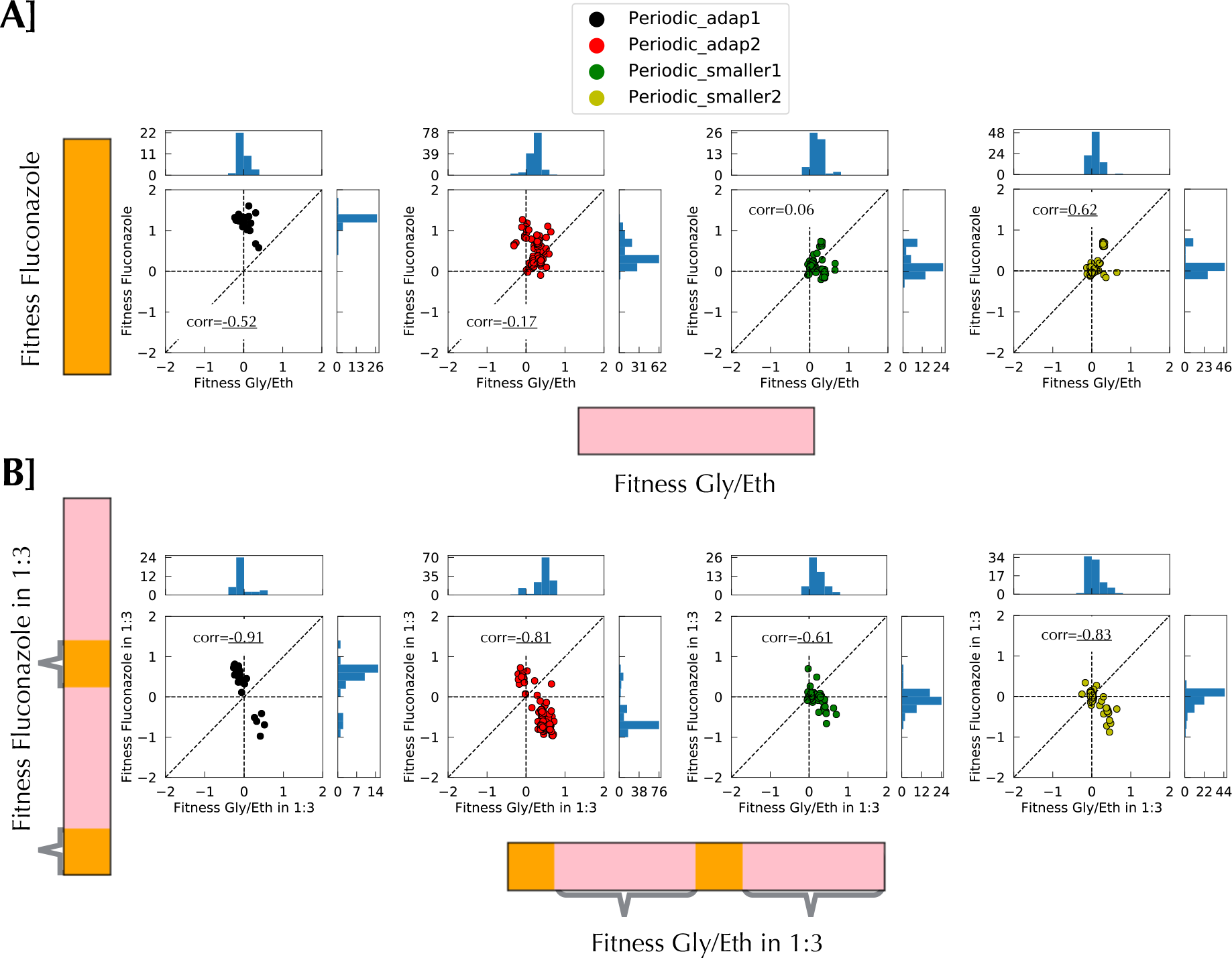
Fitness remeasurement shows different types of interaction between environments according to the time scale. **A]** Fitness measurement for Gly/Eth and Fluconazole for 40 uninterrupted generations shows no particular correlation between environments. **B]** When fitness measurement is performed on a smaller timescale in the context of switching there is a net negative correlation between the two environments. Grey braces indicate when the fitness was measured.

### The dynamics of adaptation are affected by the environmental dynamics

Visual inspection of lineage trajectories suggested that while each sequence of environments gave rise to distinct lineage dynamics (Figure 2A), some environmental sequences gave rise to similar lineage behaviors. For example, lineage trajectories from periodic_smaller1 and periodic_smaller2 are similar, because their sequences of environment are essentially the same, except they are offset from one another by a single environmental block. Likewise, periodic_adap2 and random_adap1 display visually similar lineage trajectories, likely because the lengths of the environmental blocks are similar (on the order of timescale of adaptation). By contrast, trajectories from periodic_smaller1 and periodic_smaller2 clearly differ from those of periodic_adap2 and random_adap1. To further investigate similarities and differences between conditions, we classified the environments into 3 groups: strong selection, intermediate selection and weak selection, based on the change in Shannon entropy (lineage diversity) during the evolution (Figure 2B), and both the rate at which neutral lineages went extinct and the diversity of adaptive lineages after 192 generations (Figure 2C). Under strong selection (periodic_adap1, Mix (which both initially contain Fluconazole), and random_smaller1), the diversity tends to crash early and populations are rapidly taken over by a few, fit lineages, with neutral lineages going rapidly extinct. Indeed, after 192 generations the 100 most abundant lineages are 84%, 80%, and 80% of these populations respectively. By contrast, in the weak selection environments (periodic_smaller1, periodic_smaller2) the diversity decreases around 160 generations and only a few lineages increased in frequency; after 192 generations the 100 most abundant lineages represent only 6.7% or 2.7% of the populations respectively. Under intermediate selection (periodic_adap2, random_adap1 and random_smaller2) many more lineages significantly change their frequencies, while there is still diversity in the isolated lineages: the top 100 lineages at generation 192 represent 20%, 32% and 13% of the total populations for those experiments respectively. Under weak selection (periodic_smaller1 and periodic_smaller2), diversity stays high through 192 generations and only a few lineages rise in frequency. We also used the Kolmogorov-Smirnov distance between experiments to characterize their relatedness (Figure 2D), which was largely consistent this categorization, with the exception of random_smaller2, which has a unique behavior, in that diversity didn’t decline until later, but when it did, it fell precipitously, likely driven by the emergence is a highly fit lineage that is almost fixed by 192 generations (Figure 2A).

### Environmental switching can slow down adaptation

The fitness remeasurement data allow us to better understand the differences in evolutionary behavior between different environments, how clones evolved in one environment fare in another, and how a change in environment affects fitness and adaptation (and possibly also *evolvability*). For clones isolated from any of the environments, we observe no strong deleterious fitness effects when fitness is measured in a single environment block for either of the two conditions (Fig.3A, SI11). By contrast, there is a strong negative correlation between these two conditions when fitness is measured in the context of a switching environment, such that clones often display a fitness cost in the fluconazole portion of the environment (Fig. 3B; for full data see SI 12). In Fig. 3A, fitness is measured over 40 consecutive generations in each condition separately (see SI 1), but in Fig. 3B fitness in Fluconazole is measured over 8 generations in between 24 consecutive generations in Gly/Eth, and fitness in Gly/Eth is measured following 8 generations in Fluconazole. This change of fitness behavior results in a slower rate of adaptation in periodic_smaller1 and periodic_smaller2. The deleterious effect results from the 8 generations in Fluconazole rather than the 24 generations in Gly/Eth (see SI 13,15). Indeed, fitness in the 24 generations in Gly/Eth (separated by Fluconazole) and 40 generations in Gly/Eth without switching is largely the same (SI 15). By contrast, fitness in the Fluconazole environment over 8 generations (with a switch to Gly/Eth in between) is not strongly correlated with fitness in the Fluconazole over 40 generations with no switching (SI 13). The effect of environment switching is also evident in the lineage abundances, as observable ‘zigzag’ patterns in the fitness remeasurement experiments (SI 1, bottom two panels). We hypothesize that at small timescales we are observing the effects of the switch rather than of the environments themselves; for example, the switch may lead to a change in lag phase, dependent on the new environmental block. Such a change then appears to slow down adaptation in these rapidly switching conditions.

### Environmental switching can speed up adaptation

As discussed above, sometimes environmental change can elicit a fitness cost, which can slow down adaptation, as measured by the increase in lineage abundances over time. However, in periodic_adap2 and random_smaller2, a changing environment appears to actually *increase* the rate of adaptation (Fig. 4). In both periodic_adap2 and random_smaller2, we observe little adaptation in Gly/Eth before entering the Fluconazole block, but then substantial increases in the frequencies of some lineages either at or immediately following the environment switch. These lineages show significant beneficial fitness effects in Gly/Eth (Fig. 4). It is possible that selection for lineages with modest fitness benefits in the Gly/Eth condition incidentally selects generalists that also have increased fitness in Fluconazole. The stronger selective pressure in the Fluconazole condition compared to Gly/Eth (see difference of scale in fitness SI 2 panel B) might then be the reason for this behavior, as both neutral and non-generalist lineages are then rapidly outcompeted in the face of the drug. Alternatively, some lineages may be good at switching, or instead might opportunistically take advantage of a dip in the population mean fitness due to the environmental switch.

**Figure 4:**
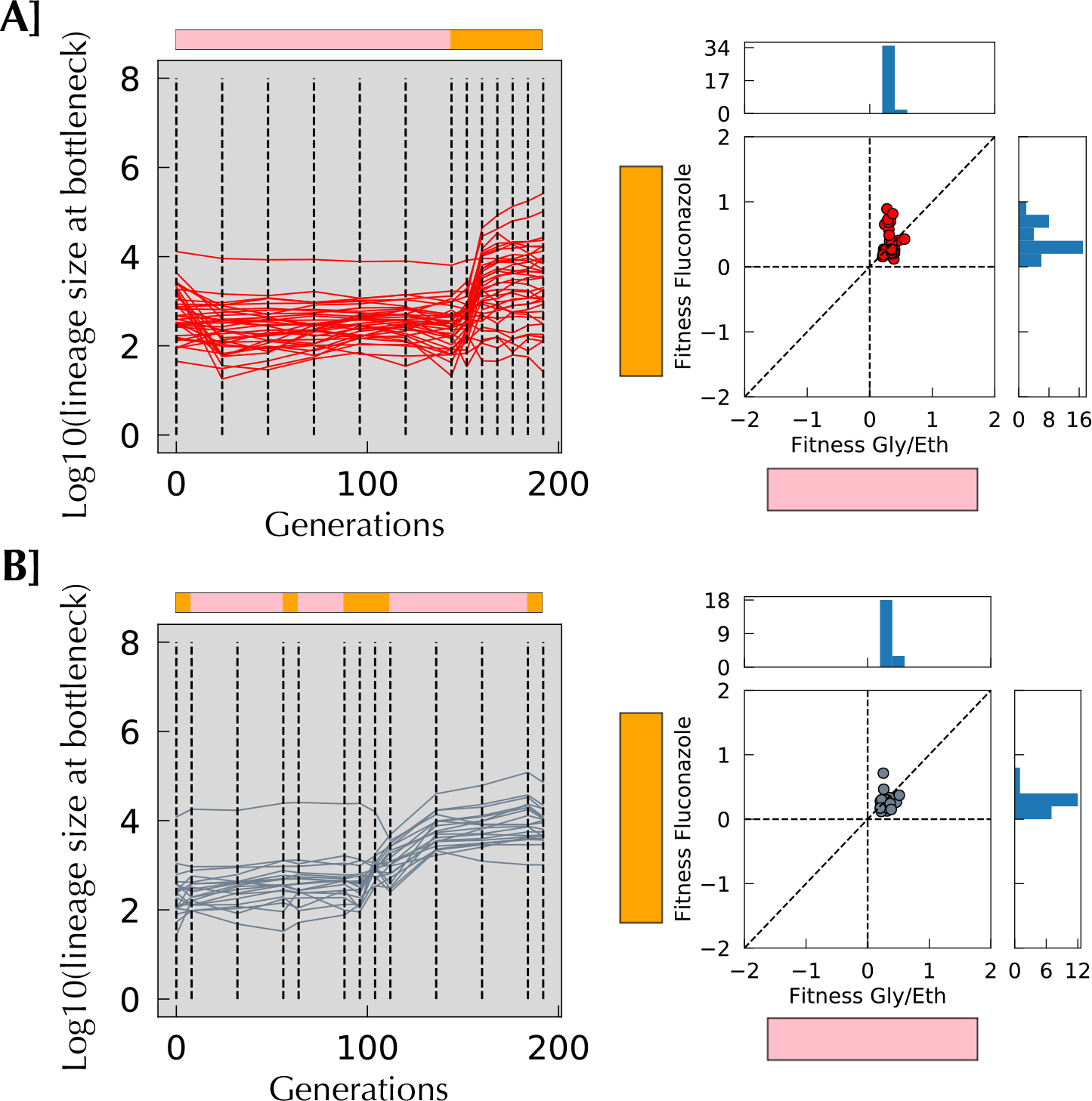
A change in the environment aids selection of lineages with high fitness in Gly/Eth. **A]** Lineages isolated from Periodic_adap2 having an average slope per cycle smaller than 0.08 in the Gly/Eth environment during the evolution yet have a remeasured fitness per cycle in Gly/Eth of > 0.2. Those fit Gly/Eth mutants were not able to reach high frequency in 144 generations in Gly/Eth, yet considerably increased their frequency during 48 generations in Fluconazole. **B]** A similar phenomenon is seen in Random_smaller2. The Fluconazole episode has reshuffled lineage frequencies: some frequent lineages decrease, while others increase. When the population goes back to Gly/Eth, a large increase in frequency can be seen.

### Different sequences of environment select for different phenotypes

To further understand how the sequence of environments affects the types of fitness benefits that are selected, we performed Principal Components Analysis on the fitness remeasurement data for all of the isolated mutants in the 5 remeasurement conditions; the first two principal components explain 89% of the variance (Fig. 5A). Based on their fitness profiles, we defined seven clusters of clones (see Methods; Fig SI 4, SI5 and 6 for threshold dependence), and examined the fitness of the clones in each cluster in each condition (Fig. 5B, 5C). Cluster 3 contains clones that are modestly more fit in all remeasurement conditions, while Cluster 5 contains clones with extreme beneficial fitness in all the remeasurement environments, suggesting clones in both clusters are generalists. By contrast, clones in cluster 7 have very high fitness in fluconazole and the mixed environment, but generally neutral fitness in Gly/Eth. Cluster 4 clones shows fitness benefits in the mixed environment, but more modest fitness gains in the switching environments (1:1, 1:3), while clones in cluster 6 show extreme fitness gains in the switching environments (1:1, 1:3), high fitness in the Gly/Eth environment, but small/average fitness in the others. Finally, cluster 1 clones only show fitness benefits in Fluconazole, with strong trade-offs in the switching environments and the Gly/Eth environment; notably, none of the other clusters showed marked trade-offs in any of the environments. Clones from a given evolving environment map to one or occasionally two clusters (Fig. 5D), while some evolving environments share some cluster usage. For example, strong initial selection for a “long” time in Fluconazole in both the Mix and periodic_adap1 environments selects for similar phenotypes in cluster 7. By contrast, a “long” time in Gly/Eth followed by a “long” time in Fluconazole may explain the similar usage of cluster 3 for clones from Periodic_adap2, Random_adap1 and random_smaller2. Finally, cluster membership for clones from both periodic_smaller1 and periodic_smaller2 is similar (clusters 3 and 4) and shares some properties with cluster membership of clones from random_smaller1, another sequence built with blocks of 8 generations in Fluconazole.

**Figure 5:**
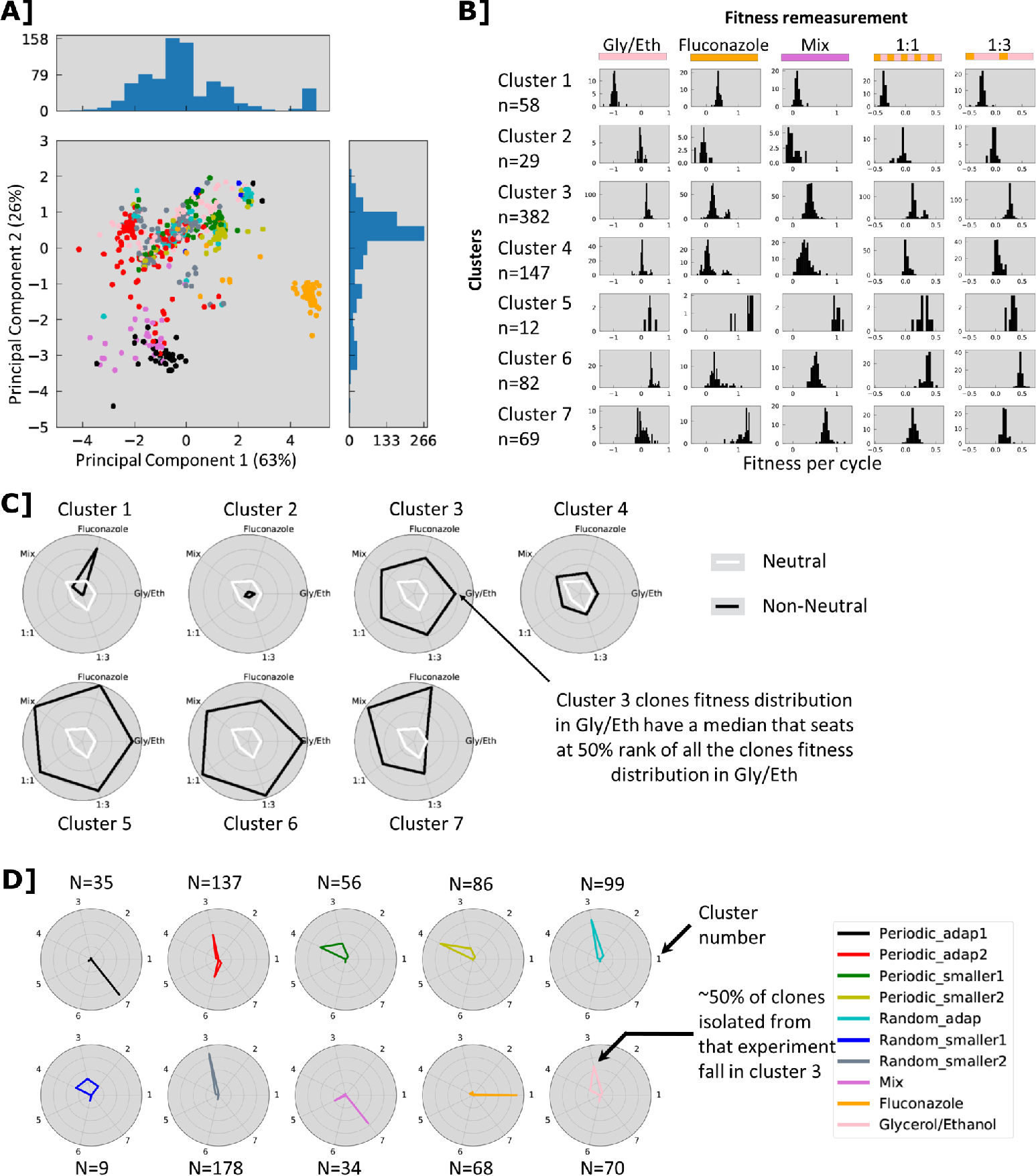
Fitness clusters. **A**] PCA analysis of combined fitness data. Each clone is represented by a five-dimensional vector of fitness values, which is projected onto a 2D space using PCA. Clones with similar fitness vectors are close together and clones from the same experiment are frequently close to one another (periodic_adap2, random_smaller2, random_adap1 or fluconazole). **B]** Distribution of fitness effects in the different remeasurement experiments for each cluster. **C]** Spider plot of cluster characteristics in the different remeasurement experiments, indicating (in black) the percentile rank of the median fitness for each cluster in a given remeasurement environment. For example, cluster 1 is highly specialized in Fluconazole whereas cluster 5 is describing generalist behavior type. White lines indicate neutral fitness in each environment. **D]** Spider plot indicating the fraction of isolated clones in each cluster from each evolution environment.

### The dynamics of the changing environment affects both the beneficial mutational spectrum and adaptive outcomes

We whole genome sequenced adaptive clones isolated from generation 192 from each evolution (7 to 51 uniquely barcoded clones per environment, for a total of 198 sequenced clones; of these, 112 had reliable fitness estimates, and 81 were considered to be non-neutral in at least one of the remeasurement conditions, i.e. adaptive); across all 198 sequenced clones, we identified a total of 482 mutations. From these, we identified genes that were recurrent targets of mutation, as they are most likely to be beneficial (Table 1). The pair of paralogous zinc finger transcription factors encoded by *PDR1* and *PDR3*, mutations in which are known to result in pleiotropic drug resistance, were frequent targets of adaptation in periodic_adap1, periodic_adap2, and Mix, likely due to selection in a “long” consecutive period in Fluconazole. Conversely, we observed frequent, heterozygous, likely loss of function mutations in *HEM3* in the periodic_smaller1 and random_smaller1 environments, which spend “short” amounts of consecutive time in Fluconazole, and for which the main fitness contribution likely comes from Gly/Eth environment. *HEM3* encodes porphobilinogen deaminase^21^, which catalyzes the third step in heme synthesis^15^. Heme is a cofactor needed for a wide variety of biological processes, including respiration and ergosterol biosynthesis; *HEM3* is essential in media lacking specific supplements and knockout mutants both lack ergosterol and fail to respire. It is unclear why decreased heme biosynthesis might be adaptive in the respiratory conditions of the Gly/Eth environment; heme is a cofactor of cytochrome C, which is responsible for the transfer of electrons between complexes III and IV in the electron transport chain. Heme is also a co-factor for cytochrome C peroxidase, which contributes to mitochondrial detoxification of hydrogen peroxide. A decreased rate of heme biosynthesis likely benefits one or both of these respiratory processes, resulting in increased fitness in the presence of a non-fermentable carbon source. Strikingly, heme is also required for sterol production (e.g. ^32^), and fluconazole itself inhibits ergosterol production, through the inhibition of the heme containing protein cytochrome P450, encoded by *ERG5*. Clustering of the fitness data for the sequenced clones shows that all but one of the *HEM3* mutants show a fitness trade-off in the fluconazole containing environments, except for the Mix environment (Figure 6). This suggests that the trade-off is not due to fluconazole itself, but instead is likely due to the switch of carbon source, and that the *HEM3* loss of function fitness benefit is specific to an environment where respiration is required.

**Table 1:** Genes mutated at least twice by 2 different non-synonymous mutations (FRS= frame_shift).

|  | Periodic_ |  |  |  | Random_ |  |  |  |
| --- | --- | --- | --- | --- | --- | --- | --- | --- |
| Gene | adap1 | adap2 | smaller1 | smaller2 | adap | smaller1 | smaller2 | Mix |
| *PDR1* | T817K, A763E, L878S, C756S<br>F769L, Q274R, G280V, P298L<br>Y270S, S753C, L1056P<br>R310W, T243A, S814Y, L537F | N234K<br>K253E |  |  |  |  |  | T817K<br>P870L<br>N234K |
| *HEM3* | L318M |  | D218E<br>K267(FRS)<br>P108L<br>G133R<br>G250(FRS) |  |  | E65K<br>A181D<br>V132M |  |  |
| *PDR3* | V219A<br>H964P | L708F |  |  |  |  |  | K272N<br>V954F<br>G948S |
| *TMN2* | P522T |  |  | V633M |  |  |  |  |
| *SGD1* | D284Y<br>M641V |  |  |  |  |  |  |  |
| *SPT23* |  | E122K | N348K |  |  |  |  |  |
| *MNE1* |  | Q284L |  |  |  |  | P121S |  |
| *LAP3* |  | D421N |  | H247Y |  |  |  |  |
| *KAP114* |  |  | I107L |  |  | I335L |  |  |
| *MGM1* |  |  | D841Y<br>L626V |  |  |  |  |  |
| *VID27* |  |  | D737A | R62K |  |  |  |  |
| *SAN1* |  |  |  | R183K<br>R185K(FRS) |  |  |  |  |
| *YHR028W-A* |  |  |  | S73C<br>S73A |  |  |  |  |
| *DYN1* |  |  | P3506S<br>A2554T |  |  |  |  |  |
| *RPO31* |  |  |  |  |  | P507A,<br>E516Q |  |  |

**Figure 6:**
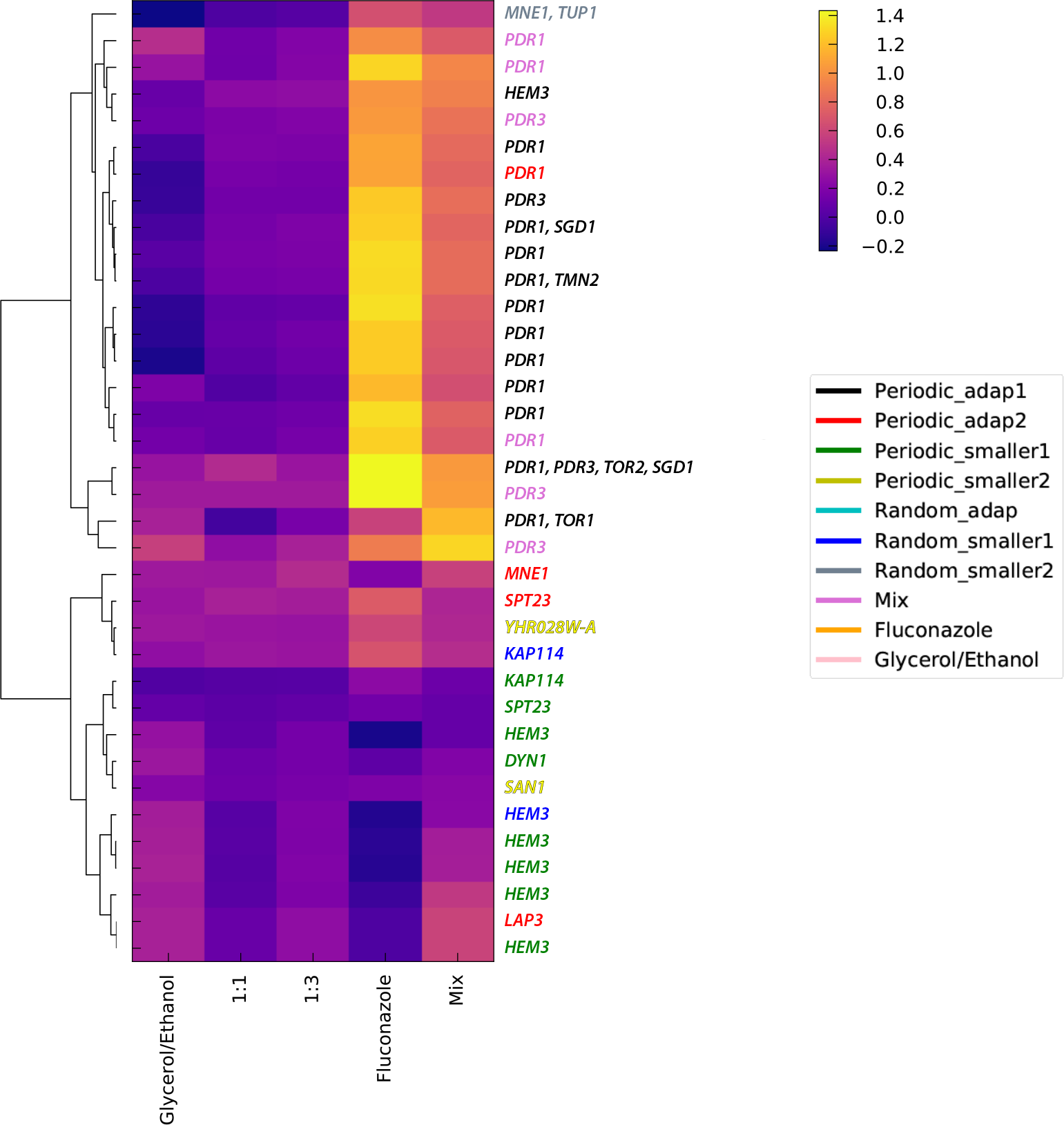
Hierarchical clustering of clone fitness data. Fitness remeasurement data, per cycle, for each clone that had reliable data in all five remeasurement conditions and at least one mutation in a gene that was recurrently mutated, were hierarchically clustered. The presumptive adaptive mutation(s) in each clone is indicated on the right, and the gene names are colored by the evolution experiment from which they were isolated.

In addition to the nature of the periodic environment influencing the beneficial mutational spectrum, it also influences the nature of adaptation itself, specifically in regard to the emergence of generalists (which have positive fitness in both growth conditions) vs. specialists (which are fit in only one or a few of the growth conditions). First, we note that selection for generalists is order dependent (Fig. 3). Indeed, strong selection for Fluconazole at the beginning of periodic_adap1, followed by selection in Gly/Eth enriched the population for lineages with high fitness in Fluconazole but approximately neutral fitness in Gly/Eth. Conversely, growth in Gly/Eth, followed by growth in the presence of Fluconazole selects for generalists, that are highly fit in Gly/Eth, with even modest fitness benefits in Fluconazole; in addition, a few mutants with high fitness in Fluconazole, of a similar magnitude to those selected in periodic_adap1, also had time to be selected (Fig. SI2 panel A, 11). This kind of generalist also arose during evolution in a consistent Gly/Eth environment (Fig. SI2, 11), despite the lack of selection in the Fluconazole environment; however, the converse is not true – clones selected in fluconazole do not show fitness gains when measured in Gly/Eth (Fig. SI2, 11), suggesting the most fit clones in Fluconazole are not generalists. Finally, a group of mutants that have high fitness in both Fluconazole and Gly/Eth arose in the Mix experiment, but those strategies were rarely observed in periodic_adap1 or 2 (Fig SI2).

## Discussion

Our data demonstrate that a population’s adaptation to a changing environment depends on the order and tempo of environmental change and the strength of selection exerted by the environment. We defined our environmental sequences using two parameters: residence time in each environment and periodicity/randomness of the switches between environments. In doing so, we shed light on how those parameters influence the outcome of adaptation in dynamic environments.

Adaptation in a varying environment will also be influenced by the joint distribution of fitness effects for adaptive mutations in each of those environments – that is, the fitness effects of all beneficial mutations from any given environment as measured across the other environment(s). If there is strong antagonistic pleiotropy between two environments, then the most fit mutations in the first environment will be strongly selected against in the second environment. The evolutionary outcome will thus depend on the time scale of adaptation relative to the switching frequency – if sufficient time is spent in the first environment for adaptive mutations to reach high frequency, the second environment is likely to subsequently select for compensatory mutations that alleviate their deleterious effects. Conversely, if a short time is spent in the first environment relative to the time scale of adaptation, the second environment will instead likely cause such mutants to go extinct. In both cases, adaptation is likely to slow down. The joint distribution of fitness effects will depend on the nature of the specific environments – correlated, or even uncorrelated environments may not greatly constrain adaptation, while anticorrelated environments will.

Our study also highlights the importance of the order of environmental conditions in determining evolutionary outcomes. We explored the simplest of ordering possibilities – with only 2 environments, one ordering is simply a shift of the alternate order. Even so, we detect a strong influence (1 then 2 or 2 then 1, i.e. periodic_adap1 and _2) at small time scales, probably driven by the difference of fitness scale between the two blocks in periodic_adap1 and 2 (Fig. SI2). The fact that we do not see any fit clones for Gly/Eth in periodic_adap1 might stem from the fact that many lineages are at high frequency after the fluconazole environment: under such conditions it becomes hard to capture the rise of mutants of small to medium fitness effect, as they are swamped by the high frequency lineages. In periodic_adap2, we observe the opposite: at the end of the first environment, Gly/Eth, lineages had not reached high frequencies, and they then encountered a new environment, Fluconazole, for which mutations with a much higher fitness effect could be selected. Nonetheless, we were unable comprehensively determine *how* environment order influences adaptation. For example, environment order may become more relevant over a longer time scale, with more switches between environments, and for a population to become adapted to a well-defined, repetitive sequence of environments, the population should likely face this repeated sequence many times. Indeed, to fully understand the influence of dynamic environments on adaptation, the time scales required may be orders of magnitude longer than needed for non-switching environments. This has two main consequences for our experiment:

First, as our experimental approach relied on having barcode diversity remaining in the population, both for lineage tracking to follow the trajectories, and fitness remeasurement (we require that lineages have different barcodes to be able to remeasure their fitness in pooled fashion), by necessity we had to focus on short-term (192 generations) evolution. Indeed, we continued the evolutions for 576 generations but one or two lineages remained. Furthermore, those lineages were already the most abundant at generation 192, meaning that lineage tracking beyond generation 192 has limited power to observe ongoing adaptation (Fig. SI31). Moreover, those lineages that fix (or nearly so) by generation 576 are already “special” by generation 192, in that they are somewhat distinct from the clusters to which they belong in the PCA projection.

Second, due to the changing environment, it is challenging to measure fitness from the lineage trajectories during the evolution itself and it is this therefore difficult to estimate the shape of the DFE. Using Maximum Likelihood inference on the lineage tracking data is less informative than when analyzing such trajectories resulting from evolution in a consistent environment. This is because in our case, four models (rather than two), have to be considered, capturing the behavior of each lineage in the different conditions as either: neutral in both, neutral then adaptive, adaptive then neutral, or adaptive then further adaptive. Distinguishing between these models is challenging, as the number of data points available to reconstruct the distribution is low. Even more challenging is the uncertainty on the identity of the remeasured clone – for example, is it representative of the lineage from which it comes? We developed an algorithm for Maximum Likelihood inference of dynamics in our changing environment data that highlight the limitations of our capacity to analyze the data that way. The power and flaws of such algorithm are depicted on simulated data (Fig. SI 17-25) and applied to our data (Fig.SI 26-30).

Both of those limitations inherent to exploring long time scales of adaptation using barcoding approaches would likely be mitigated by using an approach that allows either periodic introduction of additional barcodes (as in ^38^), or allows modification of barcodes over time (e.g. ^9^), to maintain barcode diversity within the evolving populations, and measuring fitness of isolated clones in each of the environments at each environmental switch.

## Conclusions

We characterized the impacts that dynamic environments can have on adaptation and found that switching between conditions with different dynamics can influence adaptation at multiple levels. We found that the rate of adaptation itself is influenced by switching, and that adaptation could speed up or slow down, depending on the rate of switching. When switching was fast relative to the timescale of adaptation within a condition alone, adaptation was generally slowed down, while a slower switching rate *could* speed up adaption. We also found that the order of conditions influenced the adaptive outcome: that is, conditions are not commutative, similar to the idea of priority effects in the field of ecology, such that it matters what happens first. Specifically, we found that the order could influence whether generalists were selected over specialists. Finally, different targets of adaptation were selected in different dynamic environments (even when the same amount of time had been spent in each of the different conditions), necessarily resulting in different phenotypic outcomes.

## Materials and Methods

### Yeast barcode library construction

We used 8 independently constructed barcoded diploid yeast libraries (Humphrey, Hérissant et al., in prep.) each containing ~500,000 unique barcodes. Each of these libraries bears two types of barcode: a low diversity barcode that is uniquely associated to a library and a high diversity of barcodes that is associated to a specific lineage within a library. Humphrey, Hérissant et al. first introduced the low diversity barcode as part of the landing pad (see ^26^), before the introduction of the high diversity barcode. Once the low diversity barcode was introduced, the high diversity barcode was incorporated for each strain carrying a different low diversity barcode separately.

#### Construction of the ancestor strain

Briefly Humphrey, Hérissant et al. (in prep.) generated the ancestral strain for barcoding by first crossing strain SHA321^17^, which carries the pre-landing pad, *Gal-Cre-NatMX*, at the *YBR209W* locus^26^ and the strains HR026d (see list_strains.xlsx for genotype, Mia Jaffe unpublished) which contains the Magic Marker^43^.

Mata spores derived from this cross were grown on Nourseothricin, to select for the pre-landing pad, which contains *Gal-Cre-NatMX*, and then backcrossed to FY3^47^ five times, each time selecting for NatMX and Canavanine. Spores derived from the final backcross were after one more mating with FY3 was performed to obtain the diploid ancestor (Strain GSY6699). This last cross allowed us to obtain a diploid strain heterozygous for the YBR209W locus, containing one copy of the wild type locus and one copy with the pre-landing pad.

#### Construction of barcoded landing pad strains

From the ancestor diploid strains, a low diversity barcoded landing pad was introduced. The landing pad contained lox66, an artificial intron, the 3’ half of *URA3* and *HygMX* along with the low diversity barcode.

To introduce this landing pad, Humphrey, Hérissant et al.(in prep.) amplified by PCR the fragment of interest from the plasmid library L001 (~75,000 barcodes)^17^. The PCR fragment was inserted into the genome by homologous recombination, using *NatMX* as a selectable marker. After selection using Hygromycin, the grown colonies, that were the Barcoded Landing pad strains, were isolated and saved for subsequent introduction of the high diversity library of DNA barcode.

#### Construction of high diversity libraries and final pool

Each individual diploid barcoded landing pad was then transformed using the plasmid library pBAR3-L1 (~500,000 barcodes)^26^. This plasmid carries lox71, a DNA barcode, an artificial intron, the 5’ half of *URA3*, and *HygMX*. Transformants were plated onto SC +Gal −Ura, to allow expression of the Cre recombinase, which is under the *GAL1* promoter. The recombination between lox66 and lox71 is irreversible and brings the two barcodes in close proximity to form an intron within the complete and functional *URA3* gene. Per landing pad strain, we generated between ~10,000 and 250,000 transformants. The plates were scraped, and transformants from each plate were stored separately in glycerol at −80°C.

### Experimental Evolution

The final pools, each containing ~500,000 unique barcodes, were evolved by serial batch culture in 100 mL of SC −URA media in 500 mL baffled flasks in the different sequences of environment shown in Fig.1.

In the list below, a letter represents one passage (8 generations). G stands for SC-URA 2% Ethanol/2% Glycerol (48 hours between passage), F for SC-URA 2%Glucose + 4 μg Fluconazole in 100 mL culture (24 hours between passage) and P for SC-URA 2% Ethanol/2% Glycerol 4 μg Fluconazole in 100 mL culture (48 hours between passage). All cultures were grown at 30°C; the list below corresponds to the first 192 generations.

The evolution experiments were started with a pre-culture of each pool in 100 mL of SC −URA 2% Glucose at 30°C overnight. This pre-culture was used to inoculate evolutions with ~5e7 cells (~ 400 μL).

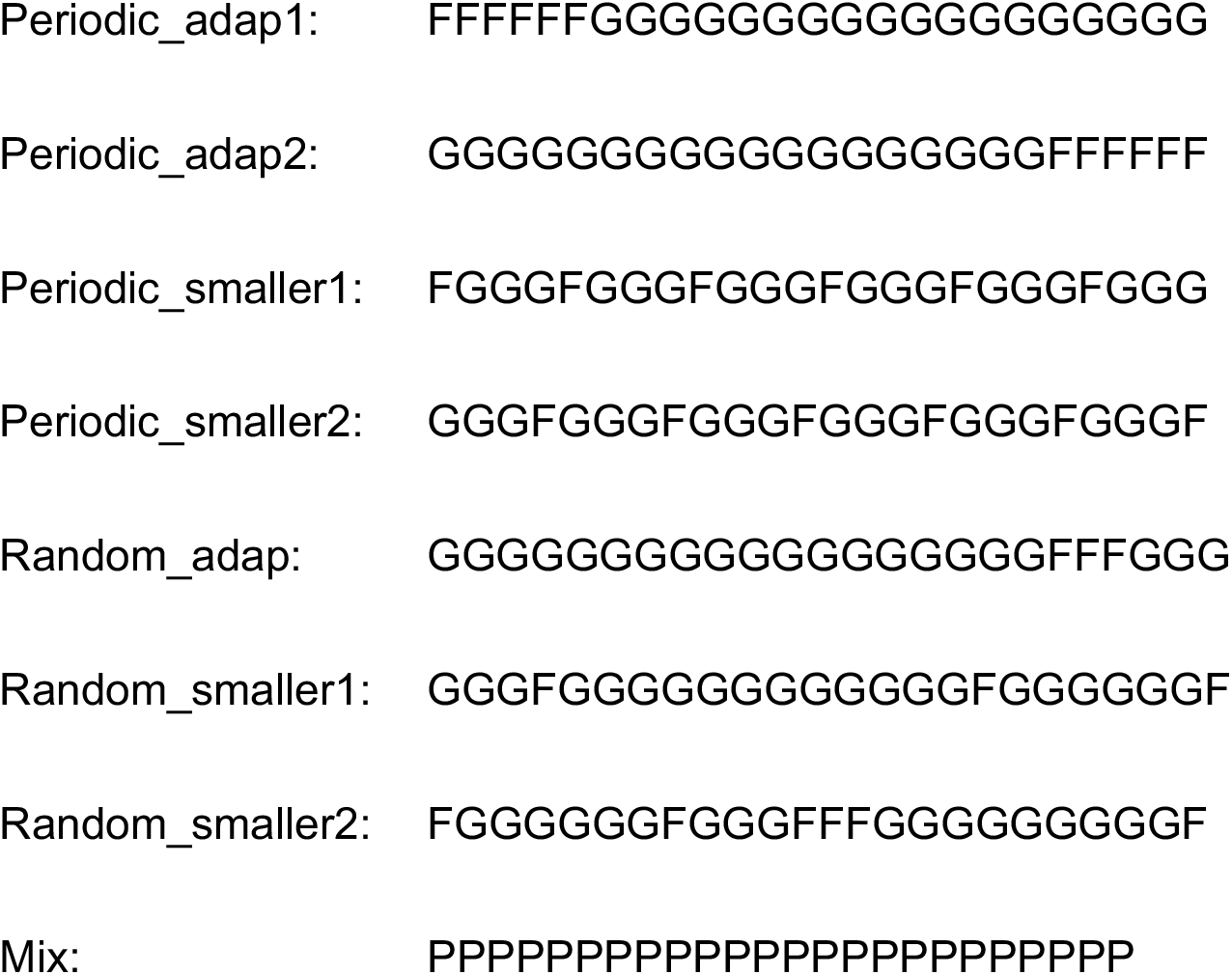

Serial transfers were performed by transferring ~5e7 cells (~ 400 μL) into fresh media. The remainder of the culture was used to make glycerol stocks; 3 tubes with 1 mL of culture each were stored at −80°C (with 16.6 % final Glycerol), while the remaining ~90 mL were centrifuged (3,000 rpm for 5 min) and resuspended in 5ml 0.9M sorbitol solution (0.9M Sorbitol, 0.1M Tris-HCL, pH7.5 0.1M EDTA, pH8.0) for storage at −20°C.

### PCR amplification of the barcode locus

#### DNA Extraction for barcode sequencing

From the sorbitol stock, DNA was extracted using the MasterPure™ Yeast DNA Purification Kit (Epicentre MPY80200), with slight modifications compared to the manufacturer’s guidelines as follows: the lysis step was performed for one hour in lysis buffer, supplemented with RNAse at 1.66 μg/μL. The DNA was washed at least twice with 70% Ethanol to remove remaining contaminants. Because the number of cells in a pellet exceeded the upper limit of the kit by roughly 6-fold, 6 extractions were performed per pellet. To do so, a cell pellet was resuspended in 900 μL of the lysis buffer and aliquoted in 6 tubes (150 μL each). The aliquots were complemented with the appropriate volume of lysis buffer (150μL, 300 μL total) to follow the manufacturer’s guidelines. We used the same procedure for DNA extraction during the lineage tracking or the fitness measurements.

#### Barcode amplification from population samples

We used a two-step PCR to amplify the barcode locus for Illumina sequencing as described^26,45^, with the following modifications. For the first step, we supplemented the PCR reaction with 2mM MgCl2 and used only 6 PCR reactions per timepoint (600 ng of genomic DNA per tube). Nevertheless, in the event of PCR failure, we performed 12 additional reactions per timepoint for the first step with the same amount of DNA, lowering the DNA concentration, each with 300 ng of genomic DNA.

The primers used for this first step are listed in list_primers.xlsx. The Ns in the primers are the Unique Molecular Identifiers (UMIs) which are random nucleotides used to uniquely tag each amplicon product for subsequent removal of PCR duplicates during downstream analysis. All primers were HPLC purified to ensure that they were the correct length.

After the first step, all tubes were pooled and purified using the QIAquick PCR Purification Kit (Qiagen, 28106). For the second step, we used Herculase II Fusion DNA Polymerase (Agilent – 600677) which is a more efficient high fidelity enzyme, with the following PCR settings: 2’ 98°C, followed by 24 cycles of (10” 98°C, 20” 69°C, 30” 68°C). The PCR reaction was performed with the standard Illumina paired-end ligation primers at recommended concentrations according to the manufacturer’s guidelines. The purified first step was split into the 3 PCR reaction tubes (15 μL each).

After amplification, the tubes were pooled and the reaction was purified using one column from QIAquick PCR Purification Kit and the DNA was eluted in 30μL of water. Finally, the eluted DNA was gel-purified from a 2% agarose gel to select the appropriate band and eliminate primer dimers using the QIAquick Gel Extraction Kit. The final gel-purified DNA was quantified using Qubit fluorometry (Life Technologies).

Samples were pooled according to their concentrations.

We used the same procedure for amplification of the barcode locus during the lineage tracking or for fitness measurements.

Barcode sequencing was performed with 2×150 paired end sequencing using NextSeq.

#### Isolation of clones and Fitness remeasurement

Samples from generations 192 and 576 were grown overnight in SC −URA and single cells were sorted into each well of 96 well plates containing 100 μL YPD using FACSJazz at the Stanford Shared FACS Facility as described previously^28^. We used four 96 wells plates per experiment. The barcodes for each well were recovered (see Barcode amplification of individual lineages in individual wells) and a single representative for each unique barcode was pooled together. We also added 96 clones/lineages that were defined as neutral from prior fitness remeasurement experiments (Humphrey, Hérissant et al., in prep.). To have a reference to what fitness type was expected in steady environment we also added 96 clones/lineages from Fluconazole (4 μg/ml) evolution in steady environment (Humphrey, Hérissant et al., in prep.) taken at generations 48 and 96 clones/lineages from a Gly/Eth evolution (2%,2%) in consistent environment taken at generation 168 (Humphrey, Hérissant et al., in prep.).

The final pool containing all barcoded clones was grown overnight in 100 mL baffled flasks in SC −URA 2% Glucose; the ancestor was grown in a separate flask. To begin the Bulk Fitness Assay, the ancestor and the pools were each mixed in a 9:1 ratio, and then ~5e7 cells were used to inoculate cultures to remeasure fitness in each of the different environments. Each fitness remeasurement was performed in triplicate. The conditions for fitness remeasurement are as follow (Fig.SI1):

- Gly/Eth: SC-URA 2% Ethanol/2% Glycerol for 40 generations with a passage at approximately every 8 generations (48 hours between passage).
- Fluconazole: SC-URA 2% Glucose + 4 μg/mL Fluconazole for 40 generations with a passage approximately every 8 generations (24 hours between passage).
- Mixture: SC-URA 2% Ethanol/2% Glycerol + 4 μg/mL Fluconazole for 40 generations with a passage approximately every 8 generations (48 hours between passage).
- 1:1: SC-URA 2% Glucose + 4 μg/mL Fluconazole for 8 generations (24 hours) then SC-URA 2% Ethanol/2% Glycerol for 8 generations (48 hours) for a total of 80 generations.
- 1:3: one passage in SC-URA 2% Glucose + 4 μg/mL Fluconazole for 8 generations (24 hours) followed by three passages in SC-URA 2% Ethanol/2% Glycerol for 8 generations (48 hours per passage) for a total of 64 generations.

#### Barcode amplification of individual lineages in individual wells

To determine the locations of the individual lineages in the 96 well plates after FACS sorting, a small volume of culture was boiled and saved. For amplification, a similar 2-step protocol was used. In the first step, each well had a unique combination of primers at a final concentration of 0.416 μM. OneTaq enzyme was used for amplification following this PCR settings: 3’ 94°C - (20” 94°C, 30” 48°C, 30” 68°C) 40 cycles. After this first step, 5μL of each well were pooled into one tube per 5 plates. After centrifuging to remove cellular debris, 20μL of the pooled mix were gel purified using QIAquick Gel Extraction Kit. The purified PCR product was then diluted 50 times for the second step of the PCR. In contrast to the previously described second step, Phusion^®^ High-Fidelity DNA Polymerase was used, following manufacturer’s instructions for 12 cycles. Finally, the PCR product was gel-purified as described above and the purified product was quantified using Qubit before mixing the different libraries.

### Whole Genome sequencing

#### DNA Extraction

The re-arrayed plates containing lineages of interest were grown in 750 mL of YPD for 2 days. DNA was extracted in 96 well format using the PureLink^®^ Pro 96 Genomic DNA Purification Kit (Thermo-K182104A). The sequencing libraries were made following the protocol previously described using Nextera technology^4,23^. We multiplexed up to 192 libraries using sets A and D primers from Nextera XT kits.

#### Analysis of whole genome sequencing data

Genome sequencing was performed with 2×150 paired end sequencing on NextSeq.

The analysis generally followed GATK best practices, as we have used previously^17,45^ (code from Humphrey, Hérissant et al.). Briefly, from the split and demultiplexed fastq files, reads were trimmed for adaptors, quality and minimum length with cutadapt 1.7.1^33^. Reads were then mapped to the yeast reference genome (*Saccharomyces_cerevisiae* R64-1-1, from SNPeff) using BWA version 0.7.10-r789^27^, variants were called with GATK’s Unified Genotyper v.3.3.0^36^ and finally the variants were annotated using SNPeff^10^. Variant filtering was performed, first by GATK recommended parameters and then using custom scripts to remove variants with low quality scores (below 150) or low coverages. Additionally, any variant in repetitive regions or low complexity regions, called using the Tandem Repeat Finder with default parameters, was excluded^6^.

## Supporting information

List of Primers

List of Reagents

List of Strains

remeasured fitness values

Lineage frequencies

Fitness remeasurement lineage data

Mutation calls for whole genome sequencing

## Acknowledgements

We wish to thank A. Agarwala, J. Blundell, D. Fisher and Dmitri Petrov for discussion. We thank J. Blundell, F. Rosenzweig and Y. Li for comments on the manuscript. We thank all members in the Sherlock lab for helpful suggestions. S.B. was supported by a Stanford Center for Computational, Human and Evolutionary Genomics (CEHG) Postdoctoral Fellowship. The work was supported by NIH grants R01 GM110275 and R35 GM131824 to G.S.

## Author Contributions

SB and GS designed the experiments. LH performed strain construction, and carried out the single environment evolutions, as well as provided advice and assistance on experiments. SB performed all other experiments and analyses. SB and GS wrote the manuscript.

