## Supplement for "Adaptation is influenced by the complexity of environmental change during evolution in a dynamic environment"

### Boyer and Sherlock Supplementary Information

#### Data analysis.

##### Hierarchical clustering of Kolmogorov Smirnov distances

Given two fitness distributions $F_k^j$ and $E_l^j$ for clones from experiments *k* and *l* and for which the fitness has been remeasured in remeasurement experiment *j*, we used the Kolmogorov Smirnov distance between those two distributions $D_{k,l}^j$, to calculate an overall distance between two experiments with $D_{k,l} = \sqrt{\sum_{j=1}^{5} {D_{k,l}^j}^2}$. Hierarchical clustering of the resulting distance matrix was performed using scipy.cluster.hierarchy module with median linkage and a max_cluster value of 4.

##### Defining Clusters from Fitness Data

Principal Components Analysis (PCA) was performed using the sklearn.decomposition module on a five dimensional space representing the fitness data from the five remeasurement experiments. The first two principal components were used to project onto a 2-dimensional space (Fig. SI 3). In this projection we identified clones that occupy the same space as our neutral clones; we then defined an ellipse around the neutral clones using their spreading along the two principal components (two times the standard deviation along PC1 and PC1) and aggregated clones within this ellipse with the neutral clones and removed them from subsequent analysis. In this resulting dataset, where each point represents a non-neutral clone (Fig.SI 4), all the pairwise Euclidian distances between clones were calculated. This distribution of distances is multimodal, which is a sign of clustering. We chose a distance of 2.5 as our threshold to define the limits of a cluster, as it is a typical intermediate distance in our data. Finally, the matrix of Euclidian distances was hierarchically clustered using the scipy.cluster.hierarchy module. Clusters were defined as subgroup of the hierarchical matrix that were separated by more than 2.5 along the diagonal. This is identical to using a maximum cluster condition of 7 in scipy.cluster.hierarchy module. Two points were initially defining their own clusters. For the sake of simplicity, we assigned them to their closest cluster neighbors. We checked for how the choice of threshold or projection affected our clusters (Fig.SI 5 and 6) and determined that our clusters are generally robust to modifying these parameters.

**Calculation of cluster characteristics**

Given a fitness distribution $A_k^j$ consisting of clones from clusters k for which the fitness has been remeasured in remeasurement experiment j, and fitness distribution $A^j$ made of clones from all the clusters together for which the fitness has been remeasured in remeasurement experiment j, we calculate the characteristic of cluster k in remeasurement experiment j by calculating its median ($median(A_k^j)$) and compare it to which quantile this fitness will represent in $A^j$.

**Fitness calculations from the fitness remeasurement experiments**

The fitness of each lineage was estimated based on the slopes of the lineage trajectories, as described previously (Venkataram et al., 2016). The software repository for the fitness estimation can be found at https://github.com/barcoding-bfa/fitness-assay-python. The code was slightly modified for fitness estimation within a sequence of environments (Fluconazole 1:1, Gly/Eth 1:1, Gly/Eth 1:3), by limiting the average of the slopes and noise parameter to the slopes and noise parameter of our interest: meaning we measure either the slopes in Gly/Eth, or the slopes in Fluconazole for the two changing environment sequences that we remeasured (1:1 and 1:3).

**Simulation of adaptation in multiple environments**

Simulations were performed as follows:

500,000 barcodes are generated. They have a mean probability of 10^-5 per generation to acquire a beneficial mutation during the 16 generations of the simulated library making process. This step is performed by drawing a random number from a Poisson distribution with mean: size_of_lineage * 1e-5. The effect of that beneficial mutation is drawn from a uniform DFE corresponding to the first environment encountered.

The sizes of lineages are calculated from a Poisson statistic centered around the size of the lineage at the previous generation multiplied by their fitness advantage. This lineage is then submitted to mutation via a non-synonymous mutation rate mu_env. The same process is used as before, i.e. a random number is drawn from a Poisson distribution with mean size_of_lineage * mu_env. If multiple mutants already exist in a lineage, the next mutations occurring can only be fed by the most frequent mutant in the lineage at the beginning of an environment. In case of multiple mutations, mutation effects are considered additive.

After 8 generations, which is the estimated time of passage, the population is rescaled to its saturation size, Ns, by drawing a random number from a Poisson distribution centered around the frequency of the lineage before passage multiplied by Ns. Finally, the population goes through a bottleneck of 1/256 size reduction, again using Poisson sampling. This is repeated through the total number of passages needed.

To limit the required memory size and computation time, all mutants that reached a size of 0 (i.e. went extinct) by the time of the environment change are erased. Thus, when the environment is changed, each mutation that has not been assigned a fitness in the new environment has a probability Pn to be neutral, Pd to be deleterious and Pb to be beneficial. Practically that happens by deciding the sign of the mutation effect due to those probabilities. An absolute value for this mutation is then drawn from a uniform DFE from which the boundaries are specific to the environment. We do not assume any relationship between the two environments for a mutation's fitness, such as correlation or anticorrelation.

Finally, for each stored time point, a sample of each lineage is draw from a Poisson distribution with mean frequency of the lineage * sequencing depth. In the presented simulations this sequencing depth per time point was 3*10^7.

**Attempted fitness estimation from lineage tracking during evolution experiments.**

We attempted to estimate fitness from the lineage tracking data generated from the evolution experiments themselves – however, the noise on these estimates is such that they were too imprecise to make any conclusions based on these estimates. Hence, we instead used fitness estimated from our fitness remeasurement experiments. However, the analysis described below is relevant for lineage tracking dynamics in dynamical environments, which was one of our initial goals, and thus we document it here for completeness. Even though the analysis on simulated data is conclusive, we cannot claim the same degree of confidence for analysis of the experimental data and thus did not include it in our claims. Codes for both for simulation and the lineage tracking analysis are available at https://github.com/SebastienBoyer/Lineage_tracking_dynamical_environment.

**Intro:** In the context of a varying environment or strong selection pressure, there are several challenges in directly applying our previous approach to estimate fitness from lineage tracking data. For example, at the start of the second environment the population is not monoclonal and is dispersed in terms of lineage sizes. In this context there is more than one model to consider for lineage fitting, instead of the usual one of "one mutant growing in a lineage in which the ancestor has a well-defined 0 fitness". Because of those different models, with different numbers of parameters, we needed a more complex criterion for model differentiation. An additional challenge comes from the estimation of the mean fitness of the population - because the population is not monoclonal, as well having a wide spread of lineage sizes, calculating mean fitness from the exponential decay of small sized lineages does not hold.

We provide a thorough study of these challenges and propose some alternatives. In addition, a jupyter notebook implementing these ideas is also provided (https://github.com/SebastienBoyer/Lineage_tracking_dynamical_environment).

**4 + 1 models.**

The mathematics underpinning the algorithm are the same as used previously (Levy, Blundell et al. 2015): we still best fit the probability distribution of the lineage sizes at time point t+1, knowing the lineage sizes at time t, and given some dynamical parameters. The difference is that we have to differentiate between 4 different models: either the lineage is only made of a neutral mutant, or of a neutral mutant and a growing beneficial mutant, or of only a non-neutral mutant (important if one wants to take into account standing variation at the beginning of the experiment or even more importantly at the change of the environment), or finally a non-neutral mutant and a growing beneficial mutant. Respectively those models need: 0 parameters, 2 parameters (s: fitness, tau: establishment time), 1 parameter (s1), or 3 parameters (s1, s, tau). The hypothesis in which only one mutant is rising in frequency within a lineage and contributes to the mean fitness of the lineage somewhat unrealistic, given the dynamic nature of selection in changing environments. Indeed, in our experiment some lineages become large (sometimes close to the same order of size as the total population) and are probably made of many different mutants, which is likely also important when the environment changes. To see the limit of these assumptions, see Fig.SI 20, 24,25 A] 4) and B] 4) for simulation and Fig.SI 26,27,28,29 panels 5 for experimental data.

We also considered one additional model, which relies on 3 parameters (s, s1 and f). However, it was less able to solve the problems encountered in fitting, and thus was not actively considered in most of those supplementary plots (Fig.SI 21 and 22). This model was called initial mix of mutants.

It considers that two mutants independently arose during the previous environment. It is a plausible hypothesis, as the rising mutant from previous environment might not have had time to fix and so share the lineage with its ancestral mutant when the environment changes. The parameters to be estimated are thus the fitness of the two original mutants (s and s1) in the new environment and their frequency in the lineage at the beginning of the environment (f). The mutant that has the highest fraction f is called original 1, the other is called original 2.

**Distribution to fit and algorithm.**

We calculate the relative frequency of the growing mutant after (n*8-tau) generations with a fitness s which is relative to the lineage fitness s1. Thus, the growing mutant fitness is always larger than the lineage fitness (if we don't do that, there is a symmetry in the log likelihood landscape that leads to nonsensical estimates) and the relative frequency of the two is straightforward to estimate. Then the average lineage size at time (n+1)*8 is just the relative abundance of the two mutants in the lineage multiplied by their exponential growth: s for the lineage mutant (original) and s+s1 for the growing mutant (mutant).

We maximize the likelihood function according to parameters {s,s1,tau} using a bounded custom implementation of the Nelder-Mead algorithm. The likelihood space can be very irregular, sometimes almost flat, and other times rugged - thus we initially randomly explore the space to have a better chance in our choice of the initial polygon. We noted that sometimes this was insufficient to give consistent results between two runs of the algorithm on the same lineage. To circumvent this problem, the algorithm is run 3 times for each lineage and the maximum of the 3 tries is used for parameter estimation.

The prior used is the same as used previously (Levy, Blundell et al. 2015), i.e. a mutation rate, a prior exponential distribution for the DFE and grid pattern to explore the parameter space. For each lineage, the algorithm is run for the 4 models followed by a nested algorithm that would privilege goodness of fit from a low number of parameters model. We explain our choice of model differentiation in more depth in separate paragraph below.

In mathematical terms, our approach is:

$$N = f(t-1) * N_e e^{\int_{t-1}^{\tau}(s1-\bar{x}(t\prime))dt\prime} \text{ (1)}$$

Equation 1 allows calculation of the lineage size N, until the establishment size of the mutant, knowing the mean fitness of the population (x_bar) and the frequency of the lineage at t-1. The estimation of the mean fitness is discussed next.

$$f_{mut}(t) = \frac{c}{s * N} e^{\int_{\tau}^{t}(s-\bar{s}(t\prime))dt\prime} \text{ (2)}$$

Equation 2 takes into account the relative component of the lineage fitness (s_bar) due to the growing mutant. Here s and s_bar are linked. Thus, to calculate (2) we proceed to simulate the dynamics that would lead to (2):

```
s_anc=0
s_mut=s
cc=1.75
delta=1
x_f=[cc/(s_mut-s_anc)*1./N]
s_bar=[x_f[0]*(s_mut-s_anc)+s_anc]
for i in range (1,t-tau):
	x_f.append(x_f[i-1]*np.exp((s_mut-s_bar[i-1])*delta))
s_bar.append(x_f[i]*(s_mut-s_anc)+s_anc)
```

$$r_{consistent}(t) = N * r_t * \frac{1}{N_e} e^{\int_{\tau}^{t}(s1+\bar{s}(t\prime)-\bar{x}(t\prime))dt\prime} \text{ (3)}$$

Equation 3 allows us to consistently (according to the s, s1 and tau hypothesis) calculate what r_t should be (the size of the lineage at time t).

$$r_{mut}(t) = f_{mut}(t) * r_{consistent}(t) \ (4)$$

$$< r_{t+1} >= r_{mut}(t) * e^{\int_t^{t+1}(s1+s-\bar{x}(t\prime))dt\prime} + (r_{consistent}(t) - r_{mut}(t)) * e^{\int_t^{t+1}(s1-\bar{x}(t\prime))dt\prime} \ (5)$$

Equation 5 is then injected to the usual P(r_t|r_t+1;s,s1,tau) as described by (1)

**Model differentiation.**

The prior for those distributions takes into account an exponential DFE with mean 0.1 fitness per generation, a mutation rate of 10^-5, and an increment of 1% for the fitness. Fitness bounds were between 0.01% and 0.2% fitness per generation for the growing mutant, and -0.1 and 0.2 for the lineage mutant (original). Establishment time bounds were [-50,30] for 48 generation in Fluconazole, [-50,130] for 144 generations in Gly/Eth and [-50,160] for 192 generations in 1:3.

The different models were ranked according to their log likelihood. If the best model needs fewer parameters than the second best, then it is taken as the best model. If it was not the case, then we employed a threshold. We checked two types of thresholds.

<u>AIC</u>

The first threshold we checked is the Aikike (modified for low number of data points) information criterion, which is a well-established approach for model differentiation. Practically, we calculate $AICc = AIC + \frac{2k^2+2k}{n-k-1}$ where k is the number of parameters of the model, n the number of point to fit the distribution, $AIC = 2k - 2\ln(L)$, and L is likelihood of the model. We then estimate the relative likelihood of the different models using the quantity exp $(\frac{AICc_{min}-AICc_j}{2})$. If the relative likelihood calculated from this formula is in favor of the smallest number of parameters model by at least 10

%, then the smallest number of parameters model is chosen over what is a better fitting model with more parameters.

The goodness of fit is often an order of magnitude worse for the fit in the second environment (Fig.S. 20 A] 3) and 15 B] 3)) and the Aikike criterion often privileged the max number of parameters. We emphasize that it might be that our model based on only one mutant rising is simply not descriptive enough, and that other macro parameters, such as the mean fitness, are not precisely estimated (see mean fitness estimation paragraphs).

<u>Differentiation criterion based on the contribution of the mutants to the lineage fitness.</u>

We thus tried to choose a model according to the proportion of lineage mean fitness that the rising mutant is expected to give. In fact, many times the Aikike criterion will choose a model for which a mutant is rising even though this mutant will represent less than 1% of the lineage mean fitness at the end of the evolution. This option for model differentiation, even though a less information theory rigorous approach, gives a more 'logical' reason to choose a threshold and so is the one that we chose to mainly use. We provide parameter estimation for both the threshold (mutant contribution to mean fitness in lineage Fig.SI 18, Aikike information criterion Fig.SI 19, for simulation) (Fig.SI 26-30, for actual experiments), but mainly explicit data analysis from the mean fitness of lineage criterion.

**Mean fitness estimation.**

For calculating the mean fitness, we cannot use directly what was used in (Levy, Blundell et al. 2015). In many experiments (mainly after the change of environment but also sometimes for the first environment i.e. periodic_adap1) the populations are not isogenic and are broadly distributed in terms of lineage sizes.

<u>Mean fitness from a beam of lineages behaving similarly: thread method.</u>

To solve this problem, we estimate the mean fitness from the slopes of the lineage tracking. From this estimation we focus on a subsample of lineages that behave with the same fitness and those lineages become our new ancestor and reference for fitness. We call those lineages a thread. It is then normal to see an offset between measured fitness and estimated fitness. We then use the approach from (Levy, Blundell et al. 2015) to get an estimate of the mean fitness and kappa for that subset of lineages that behave the same way, again from the same Maximum Likelihood estimation of probability of lineage size a t+1 knowing lineage size at t (Fig.SI 17).

This is a crude estimation of how the mean fitness behaves, which can lead to a poor absolute likelihood. It renders the estimation of the dynamical parameters less precise. Indeed if the lineages that are used are among the already high fitness lineages in the population ,then they would be largely insensitive to changes of mean fitness that are below their own fitness: i.e. we lack a good part of the coarse grained dynamics of the population for several a few time point.

<u>Population mean fitness calculated from known or remeasured lineage fitness.</u>

To determine whether this was a reasonable approach for estimating the dynamics of the most frequent lineages in the evolution, we calculated the mean fitness of our population in simulation and for second environment of our experiment, using the lineages' known or remeasured fitness. There is no evidence that doing so for the first environment would be of any use, since fitness is remeasured at the end of the second environment. To do so, we use exactly the same P(n(t+1)|n(t)) but this time with x_bar(t) (the population mean fitness) being the parameter to estimate. We focus on lineages that (visually for the experimental data) behave as they were having this fitness from the beginning (i.e. no second mutation rising). All the approaches to calculate fitness are compared in Fig.SI 17. Also, lineages fit from those different mean fitnesses are compared (Fig.SI 24 and 25) for simulation and the experimental data. Except for the distribution of goodness of fit, which is orders of magnitude better with the mean fitness calculated from known lineage fitness, the fitness estimations are not much changed.

**Conclusion:** We emphasize that the best lineages fit are from a mean fitness calculated from known lineages' fitness: we strongly recommend that those reference lineage fitness should be remeasured for any future lineage tracking in a fluctuating environment (or even in steady environment). We still lack good information theory on the way to choose the right model. Yet our criterion on the lineage mean fitness contribution of the rising mutant makes sense. In terms of distribution, the different parameters are differently affected by the different model choice criterion and population mean fitness estimation. Getting the fitness of the original mutant in the lineage (not the fitness of the rising mutant) is robust. But for the rising mutant and to a similar extent for the establishment times it is not as robust. This is mainly due to flatness of the Log likelihood function maximum for those parameters and so is more due to information content in the data than the algorithm itself.

In conclusion we have found a convincing algorithm to analyze lineage tracking data when environment is changing or when population is not monoclonal. It works to a great extent for dynamical parameters estimation in our simulation. Finally, when applied to our experimental data what we seem to lack is more an extended understanding of the dynamic in Fluconazole environment rather than a problem related to change of selective pressure or standing genetic variation.

**Supplementary Figures**

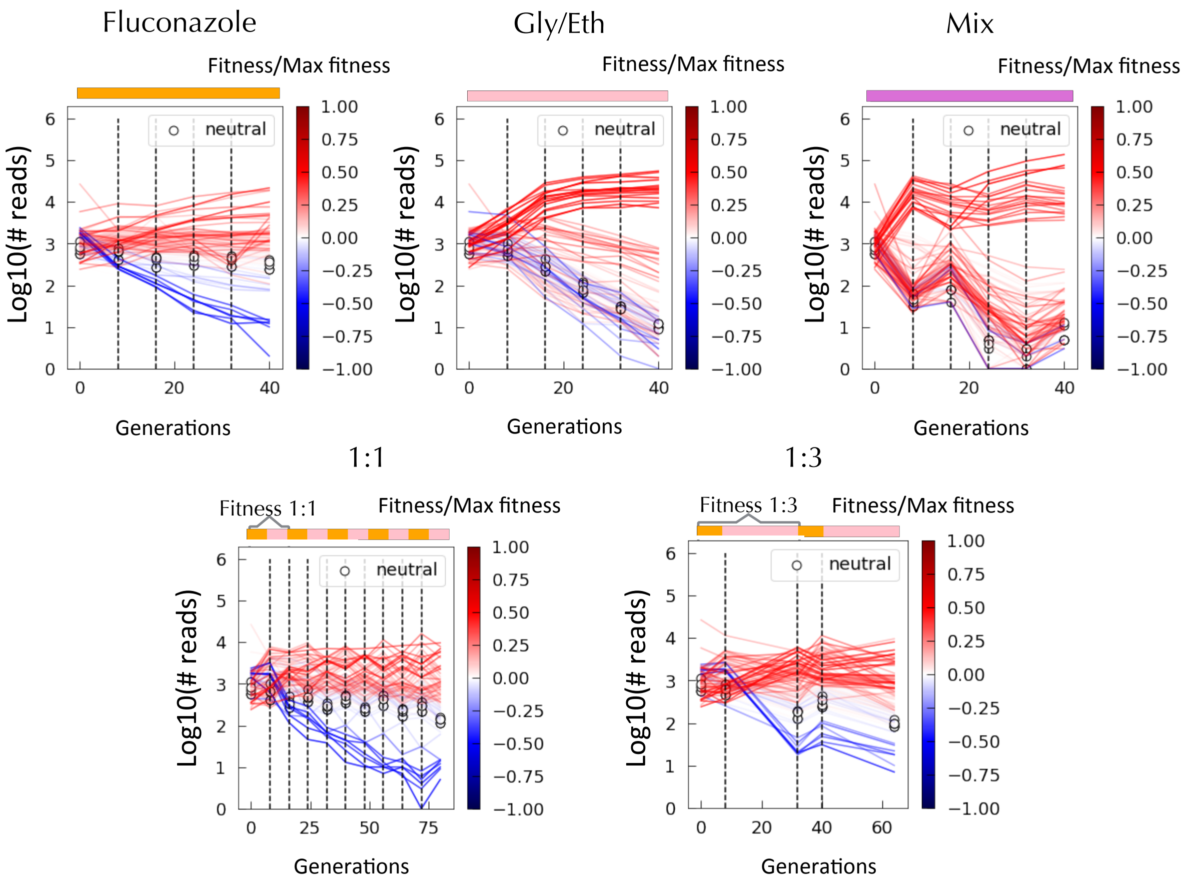


**Figure SI 1: Fitness remeasurement.** Each plot represents one of three replicates for lineage tracking fitness remeasurement experiments in the 5 different conditions. Dashed lines represent measured time points and the upper color strip indicates the environment. For condition 1:1 and 1:3, the line above the color strip indicates over which interval of environment the fitness is remeasured. Lineages known to be neutral (added in the remeasured population for that purpose) are depicted by a hollow circles. Lineages are color-coded according to their fitness proximity to either the most fit or deleterious lineage: negative fitness mutants are compared to maximally deleterious mutant and positive fitness mutants to the fittest mutants.

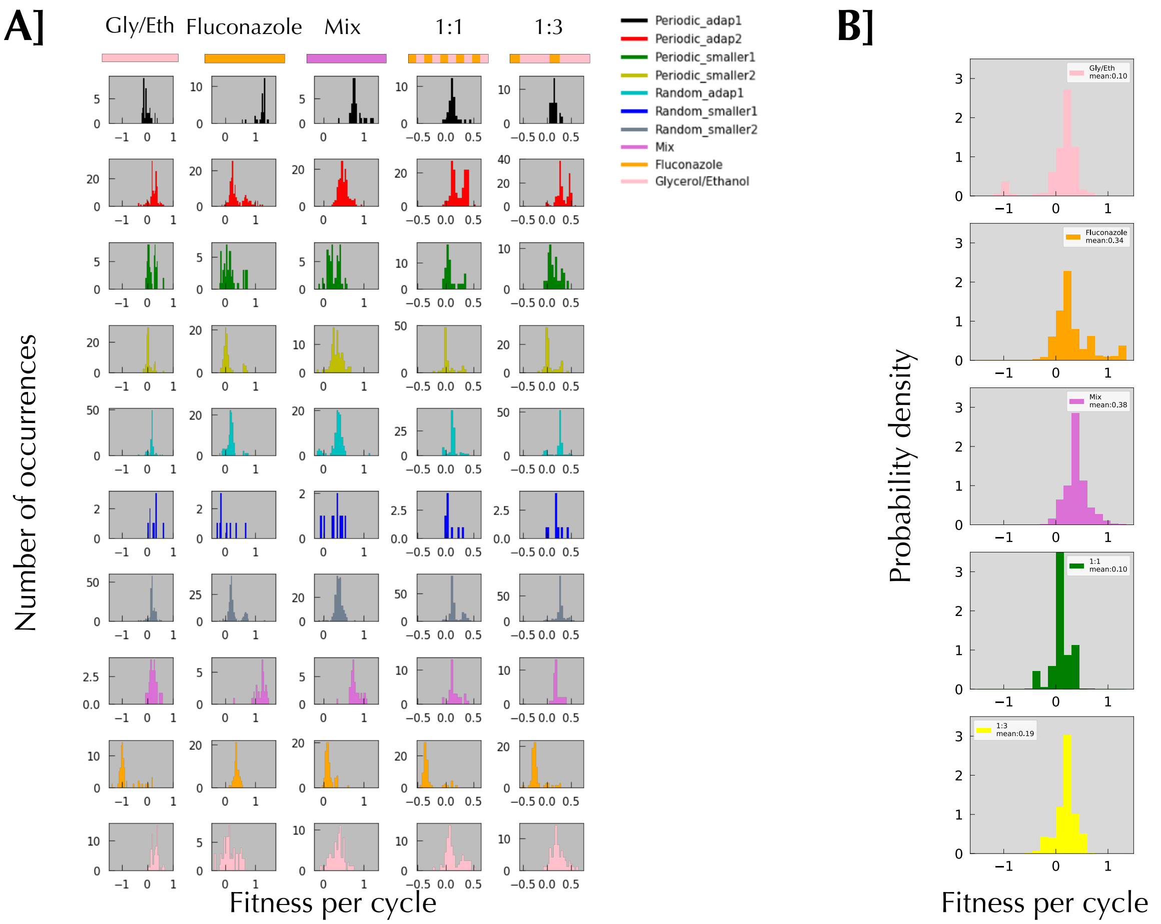


**Figure SI 2: Fitness remeasurement distributions in different conditions for non-neutral clones according to the experiment they are from. A]** Rows correspond to a specific dynamic environment from which the clones were picked. Columns correspond to the fitness distribution of those clones in the 5 remeasurement experiments. **B]** Fitness probability density distribution for the 5 different remeasurement experiments. The mean of the fitness distribution is given for each of the remeasurement experiments.

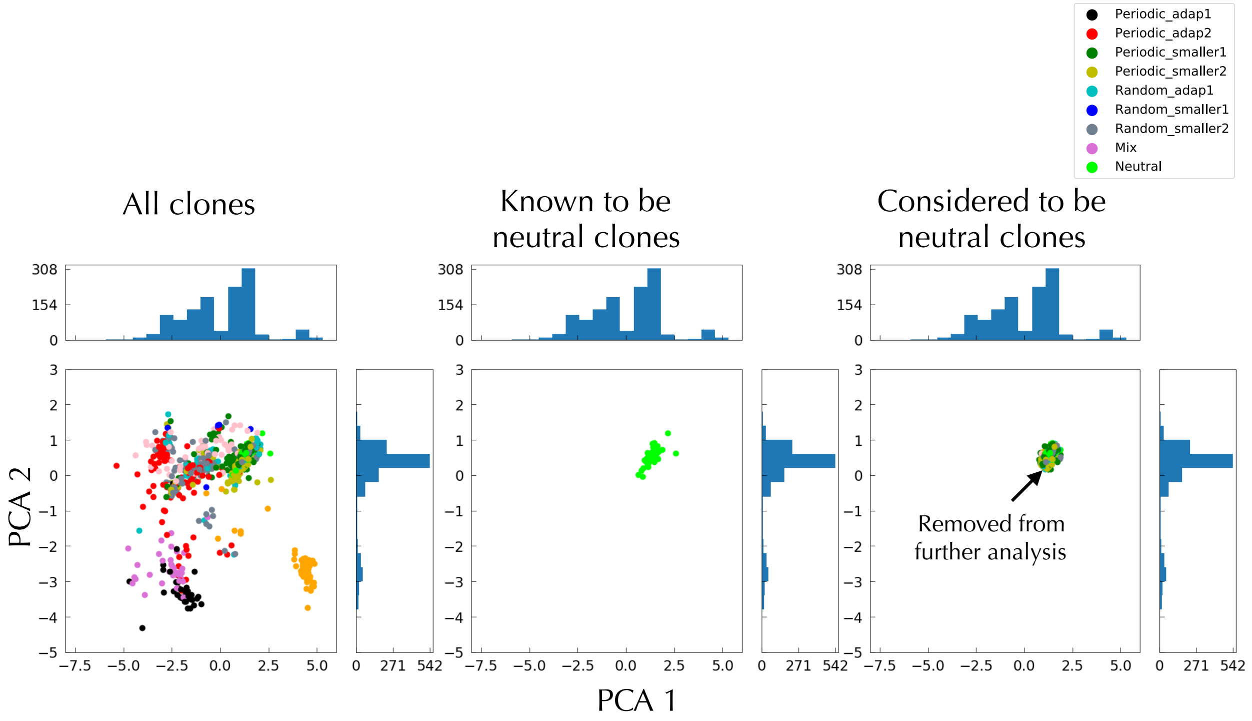


**Figure SI 3: Definition and removal of neutral clones.** Each clone is represented by a 5 dimensional vector containing their fitness in the 5 fitness remeasurement environments. We then project that 5 dimensional space on a 2 dimensional space along the first two principal components. Clones known to be neutral are grouped together forming what can be approximated by an ellipse. This ellipse is defined by its spreading (twice the standard deviation) along the PC1 and PC2 axes. Clones falling inside that ellipse are considered to be neutral.

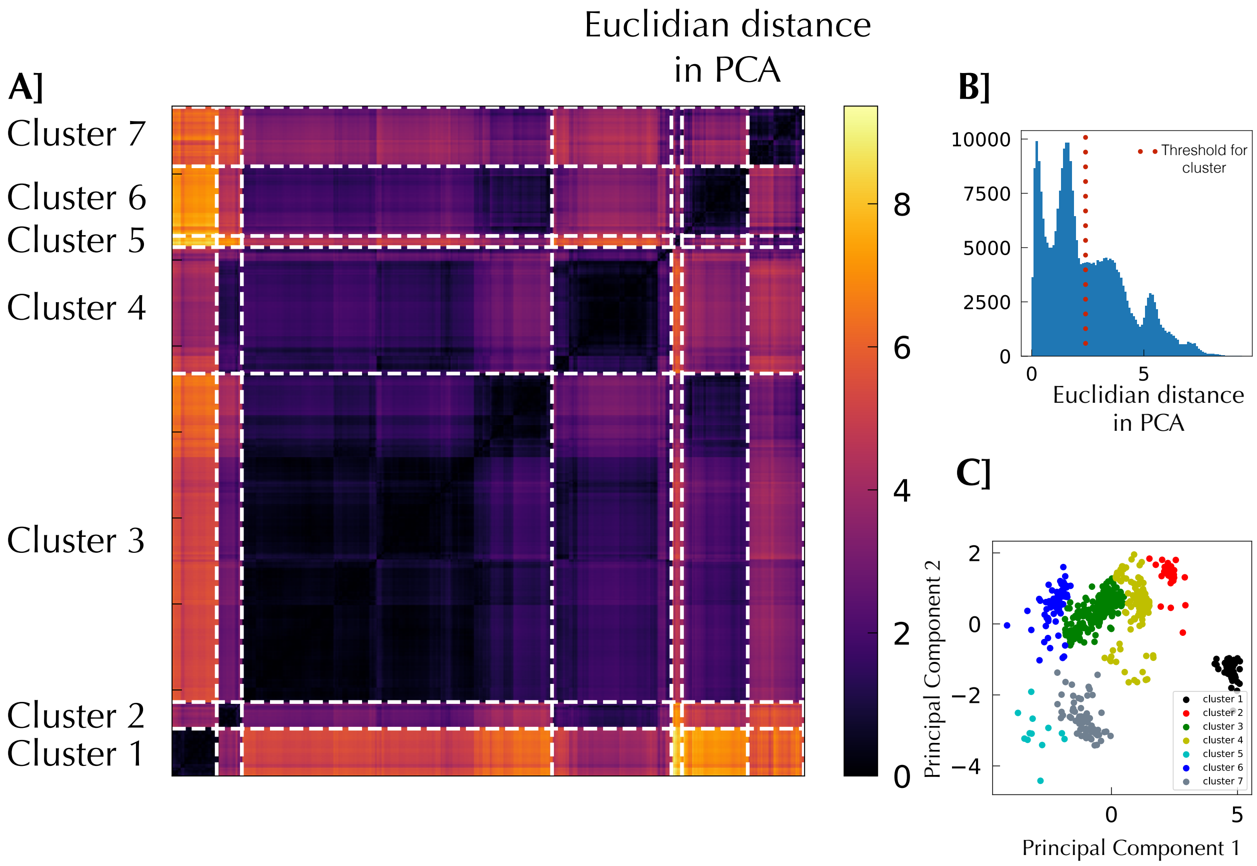


**Figure SI 4: Cluster definition from Euclidian distance in the 2D PCA projection with threshold 2.5. A] Hierarchical cluster matrix of Euclidian distances between clones in the PCA projection.** White dashed lines delimit clusters, defined by choosing a distance as defined below. **B] Distribution of Euclidian distances between clones in the PCA projection.** We chose a threshold of 2.5 to separate clusters. **C] Cluster coloring in the PCA projection.** Based on our clustering, clones are colored according to which cluster they fall in.

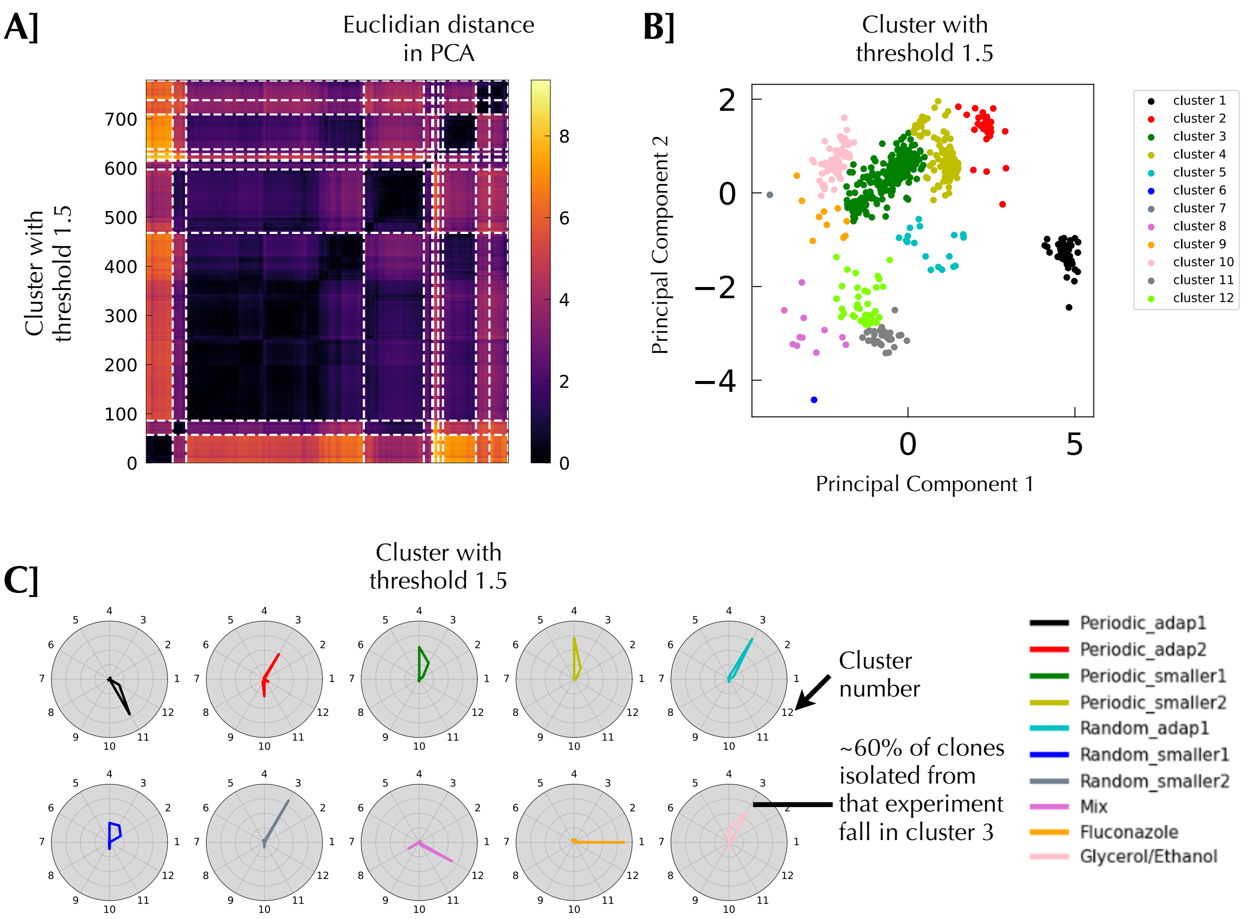


**Figure SI 5: Effect of distance threshold on clustering A] Hierarchical cluster matrix of Euclidian distance between clones in the PCA projection, using a threshold of 1.5. B] Cluster coloring in the PCA. C] Spider plots of cluster repartitioning with different threshold.** These spider plots are very similar to **Fig4.C** and shows that the choice of the threshold is not very critical to our conclusion and fitness clustering: periodic smaller 1 and 2 still show the same usage of clusters as do periodic adap2 random adap1 random smaller2 and Glycerol/Ethanol.

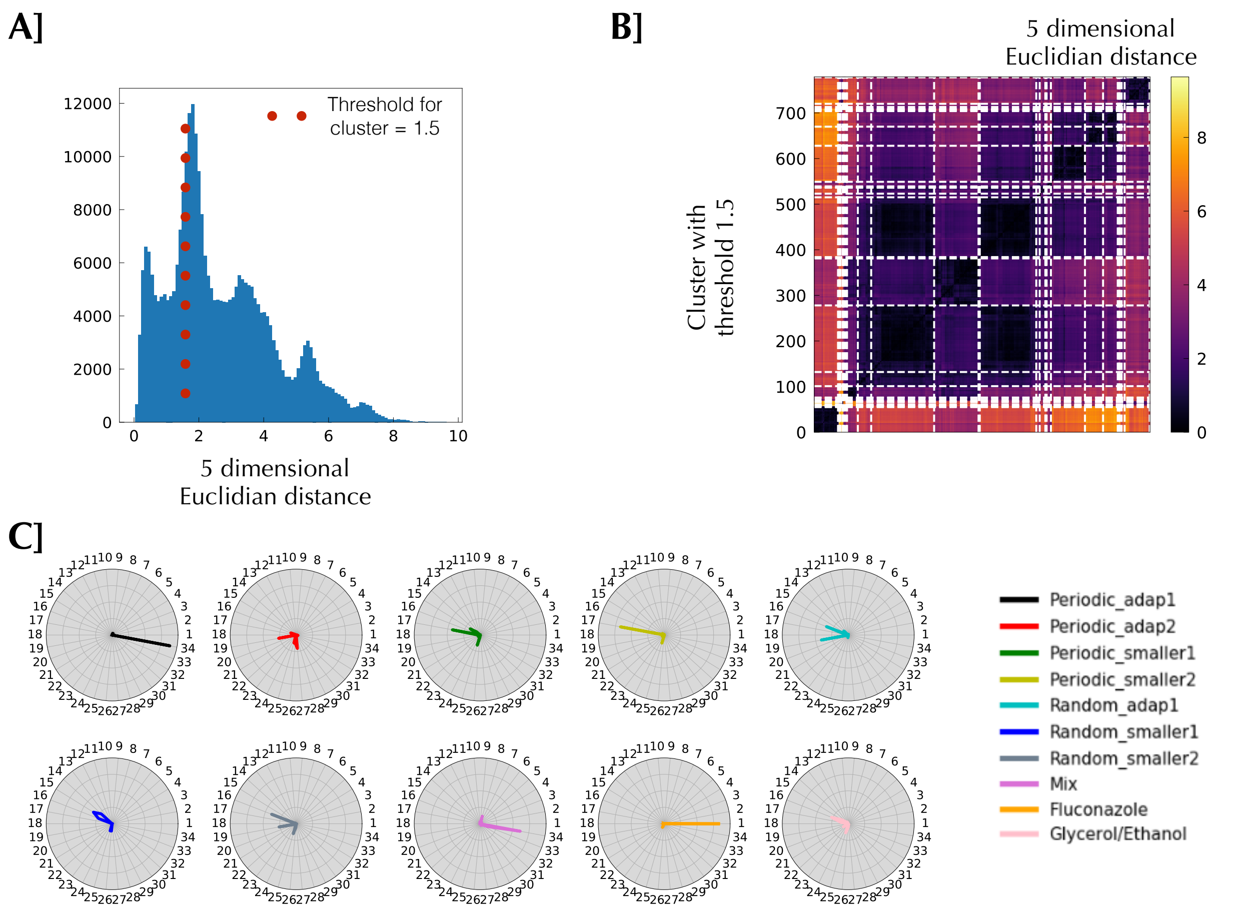


**Figure SI 6: Clusters from the 5 dimensional Euclidian distances. A] Distribution of Euclidian distances between clones in the 5 dimensional space.** To calculate those distances, the fitness measurements were rescaled by their standard deviation so that they have the same weight. Here again we see different length scales and chose 1.5 as a threshold between clusters. **B] Hierarchical cluster matrix of Euclidian distance between clones in the 5 dimensional space.** With the 1.5 threshold we have 34 clusters. **C] Spider plots of clusters repartition in our evolution experiment.** The threshold choice, or even projection into a lower dimensional space does not affect our conclusions.

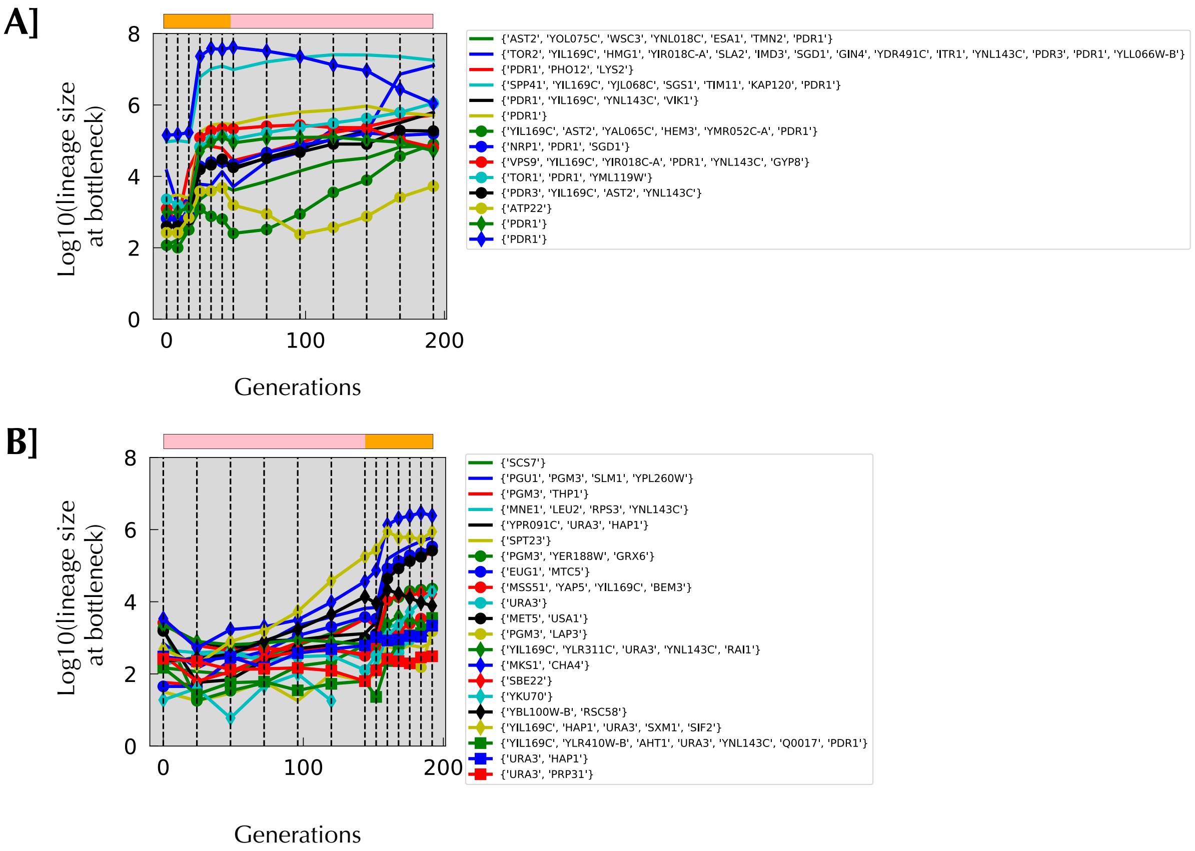


**Figure SI 7: Lineage tracking data for whole genome sequenced clones and their associated mutated genes, for A] periodic_adap1 and B] periodic_adap2.** Mutated genes that passed the different confidence thresholds are shown; for genes that appear multiple times which are not in table 1, the **same exact** mutation was observed multiple times.

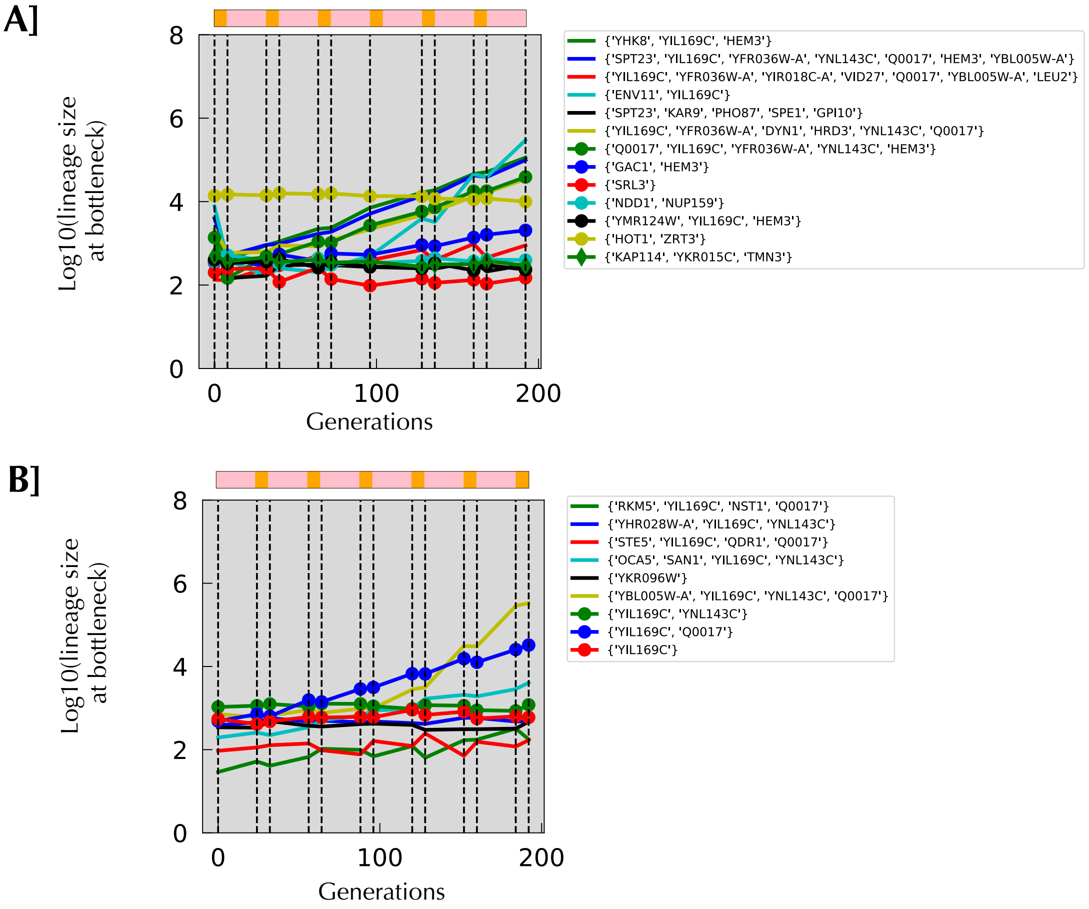


**Figure SI 8: Lineage tracking data for whole genome sequenced clones and their associated mutated genes, for A] periodic_smaller1 and B] periodic_smaller2.**

A]

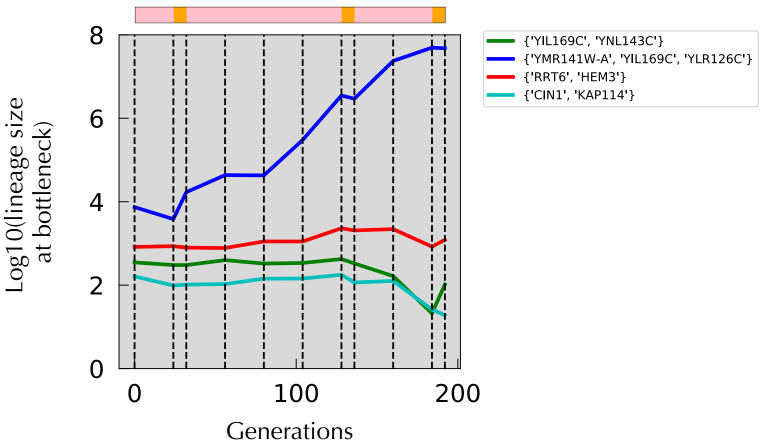


B]

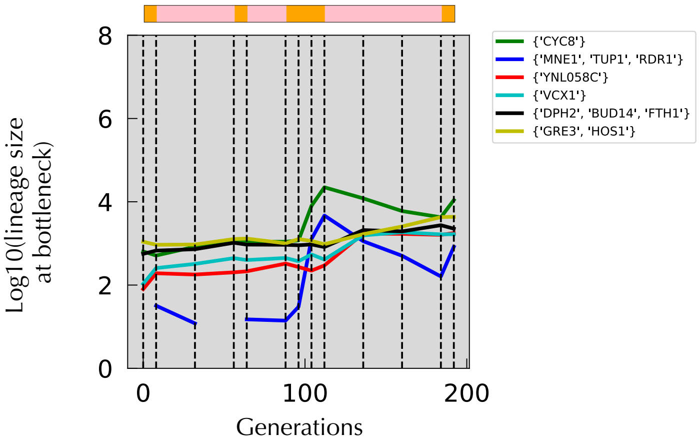


**Figure SI 9: Lineage tracking data for whole genome sequenced clones and their associated mutated genes, for A] random_smaller1 and B] random_smaller2.**

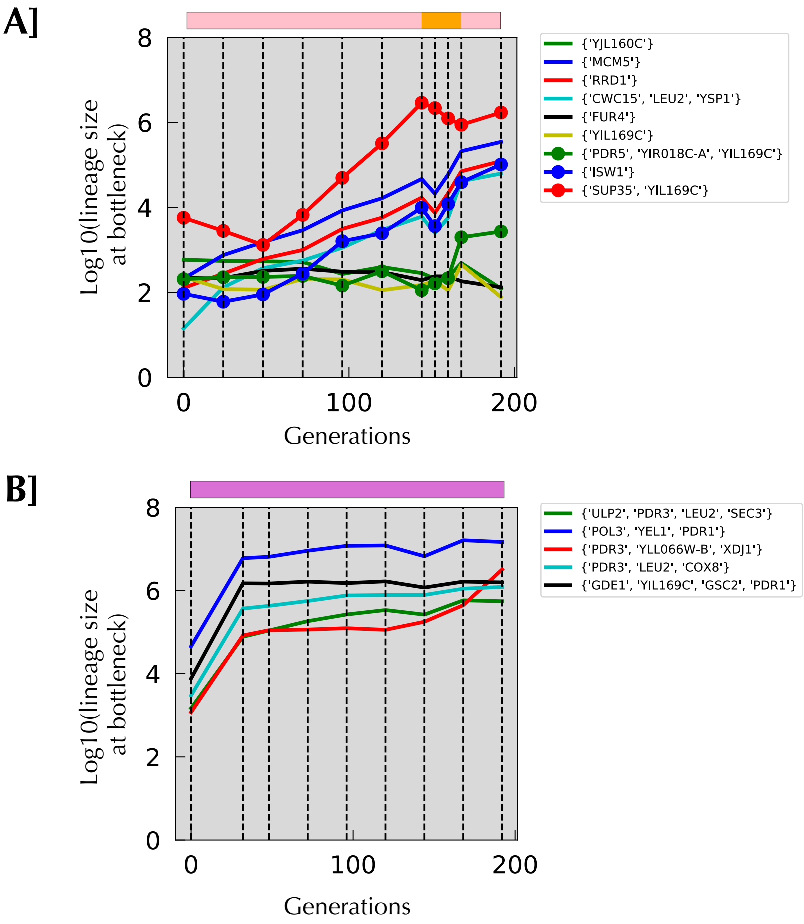


**Figure SI 10: Lineage tracking data for whole genome sequenced clones and their associated mutated genes, for A] random_adap and B] mix.**

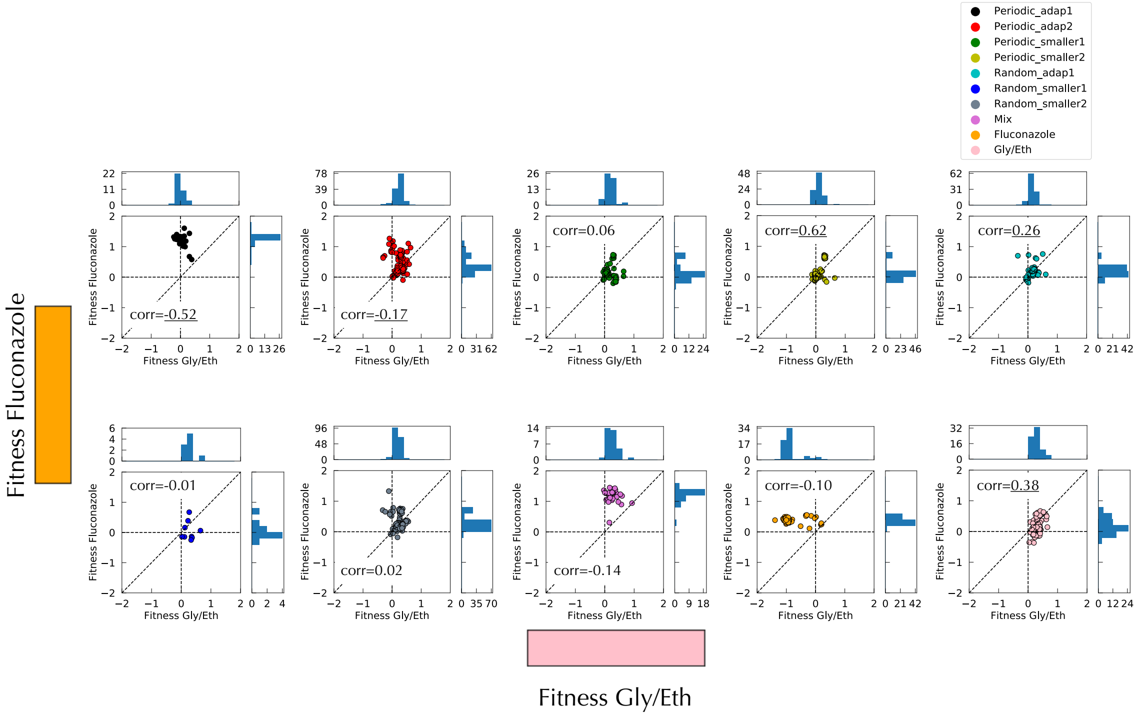


**Figure SI 11: Relationship between fitness per cycle in Fluconazole and fitness per cycle in Gly/Eth.** Underlined correlation indicates a P-value<0.05.

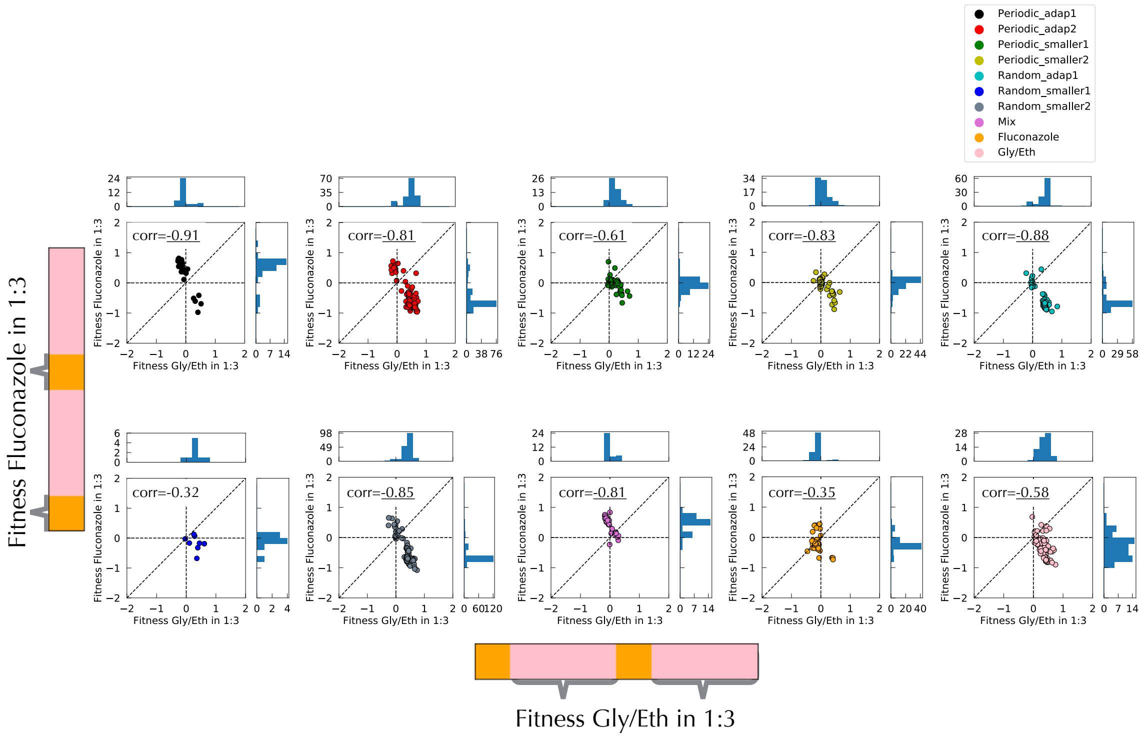


**Figure SI 12: Relationship between fitness per cycle in Fluconazole 1:3 and fitness per cycle in Gly/Eth 1:3.** Underlined correlation indicates a P-value<0.05. Braces on the color strip indicate in which block of environment fitness was remeasured.

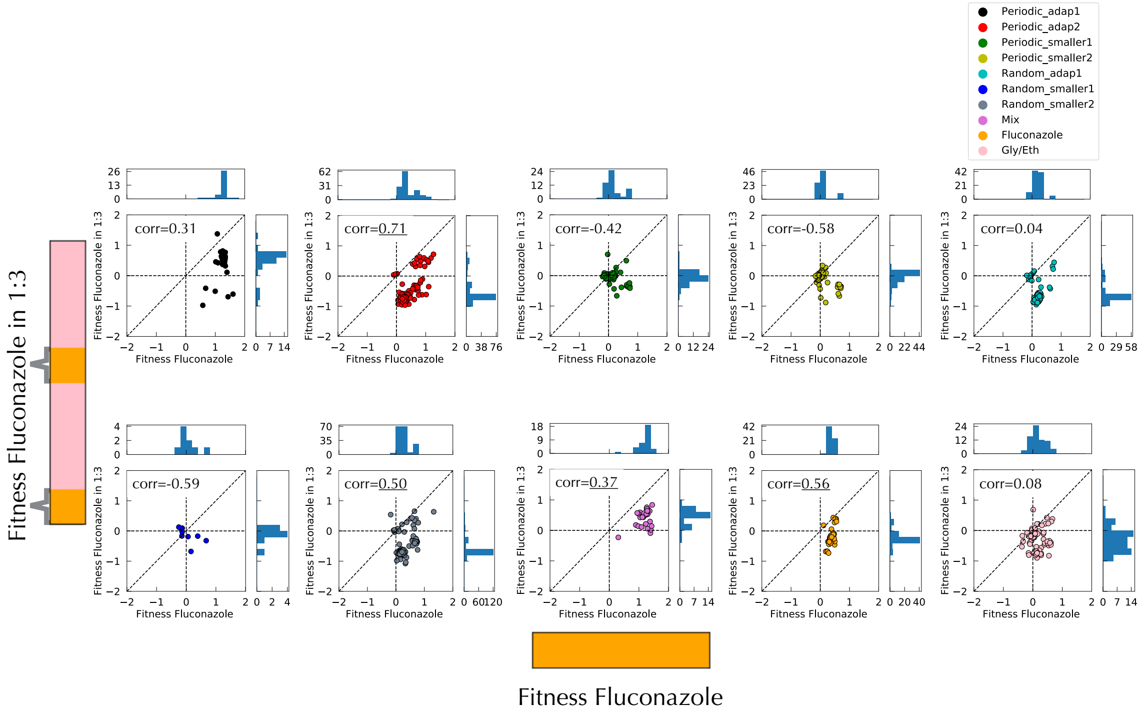


**Figure SI 13: Relationship between fitness per cycle in Fluconazole 1:3 and fitness per cycle in Fluconazole.** Underlined correlation indicates a P-value<0.05. Braces on the color strip indicate in which block of environment fitness was remeasured.

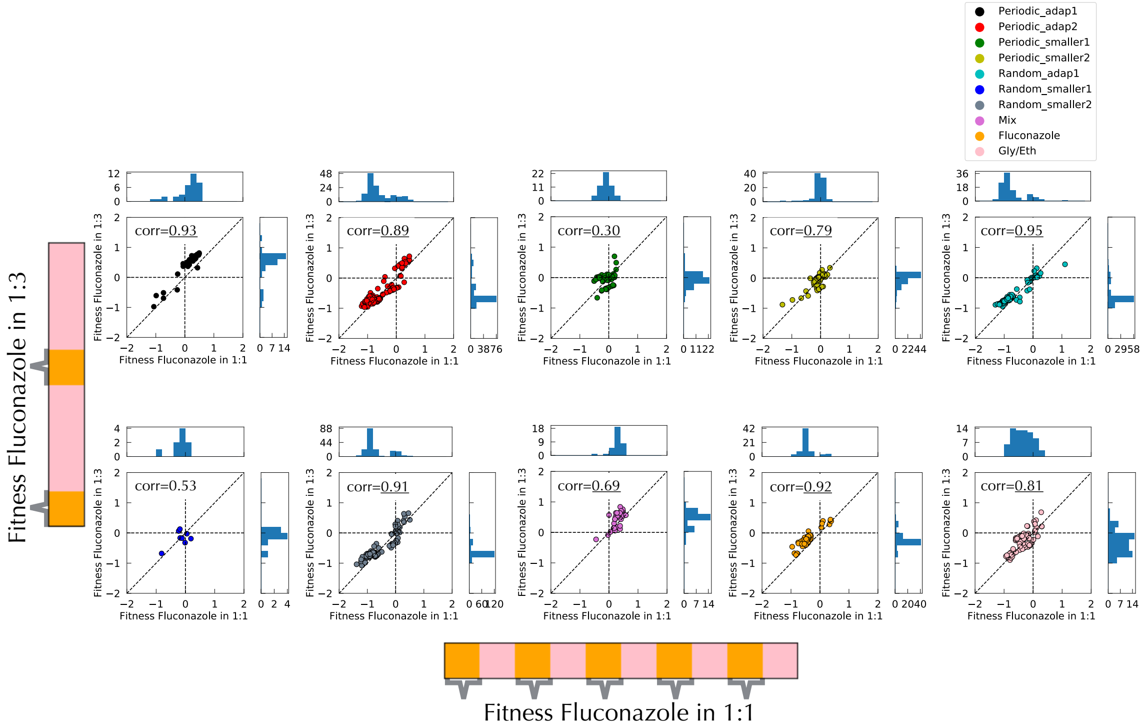


**Figure SI 14: Relationship between fitness per cycle in Fluconazole 1:1 and fitness per cycle in Fluconazole 1:3.** Underlined correlation indicates a correlation with P-value<0.05. Braces on the color strip indicate in which block of environment fitness was remeasured.

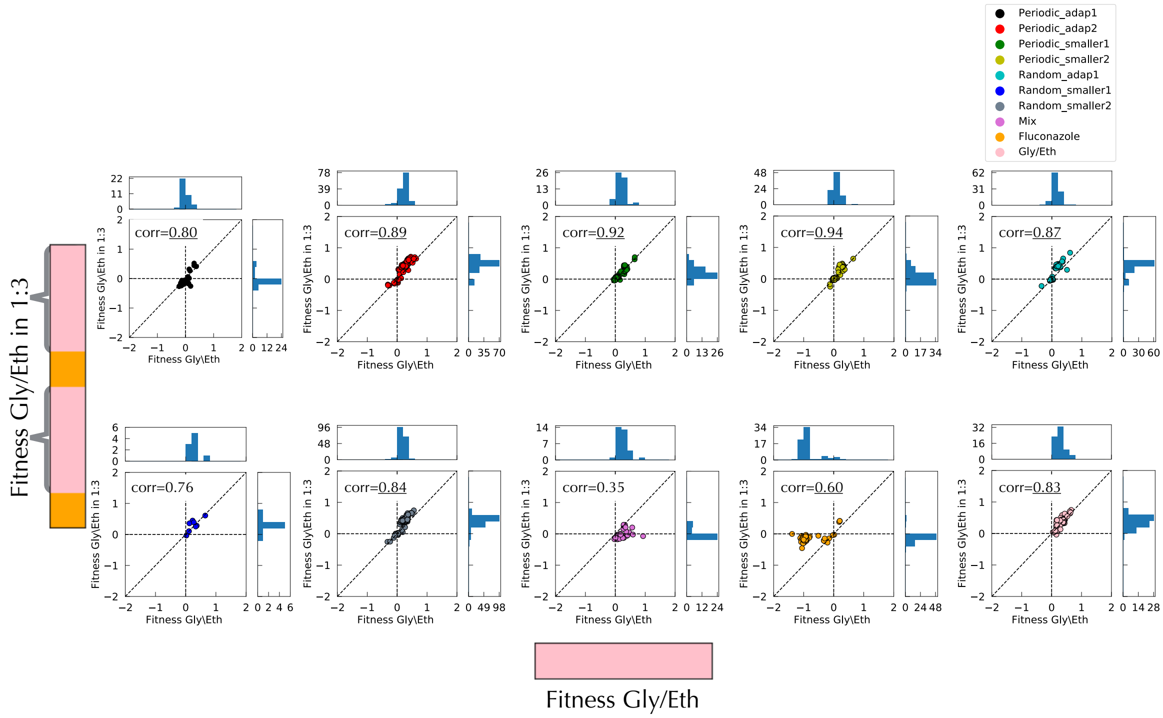


**Figure SI 15: Relationship between fitness per cycle in Gly/Eth 1:3 and fitness per cycle in Gly/Eth.** Underlined correlation indicates a P-value<0.05. Braces on the color strip indicate in which block of environment fitness was remeasured.

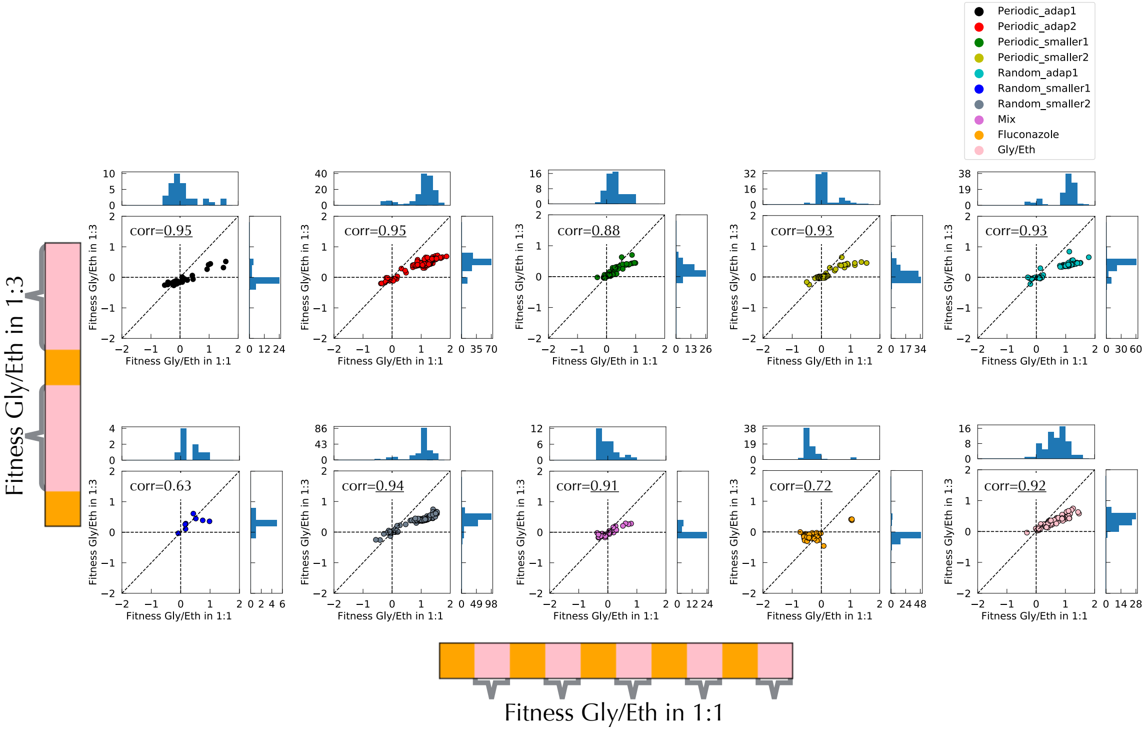


**Figure SI 16: Relationship between fitness per cycle in Gly/Eth 1:3 and fitness per cycle in in Gly/Eth.** Underlined correlation indicates a P-value<0.05.

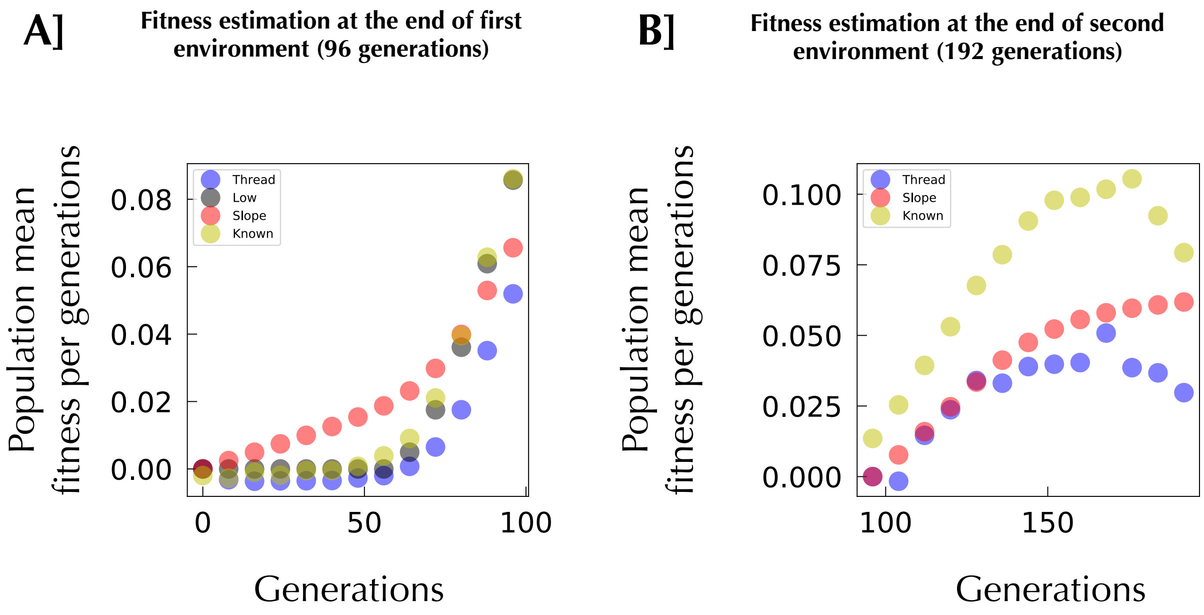


**Figure SI 17 Simulation: population mean fitness estimation by different mean. A] Population mean fitness evolution in simulation: first environment.** In blue the mean fitness is calculated using a thread of similarly behaving lineages, as described in SI. In red, the mean fitness is calculated from the slopes of the lineage tracking data. In black, mean fitness is derived from exponentially decaying small size lineages, as described (Levy et al, 2015). In yellow mean fitness is fitted using lineages for which we know the fitness. **B] Mean fitness evolution in simulation: second environment.**

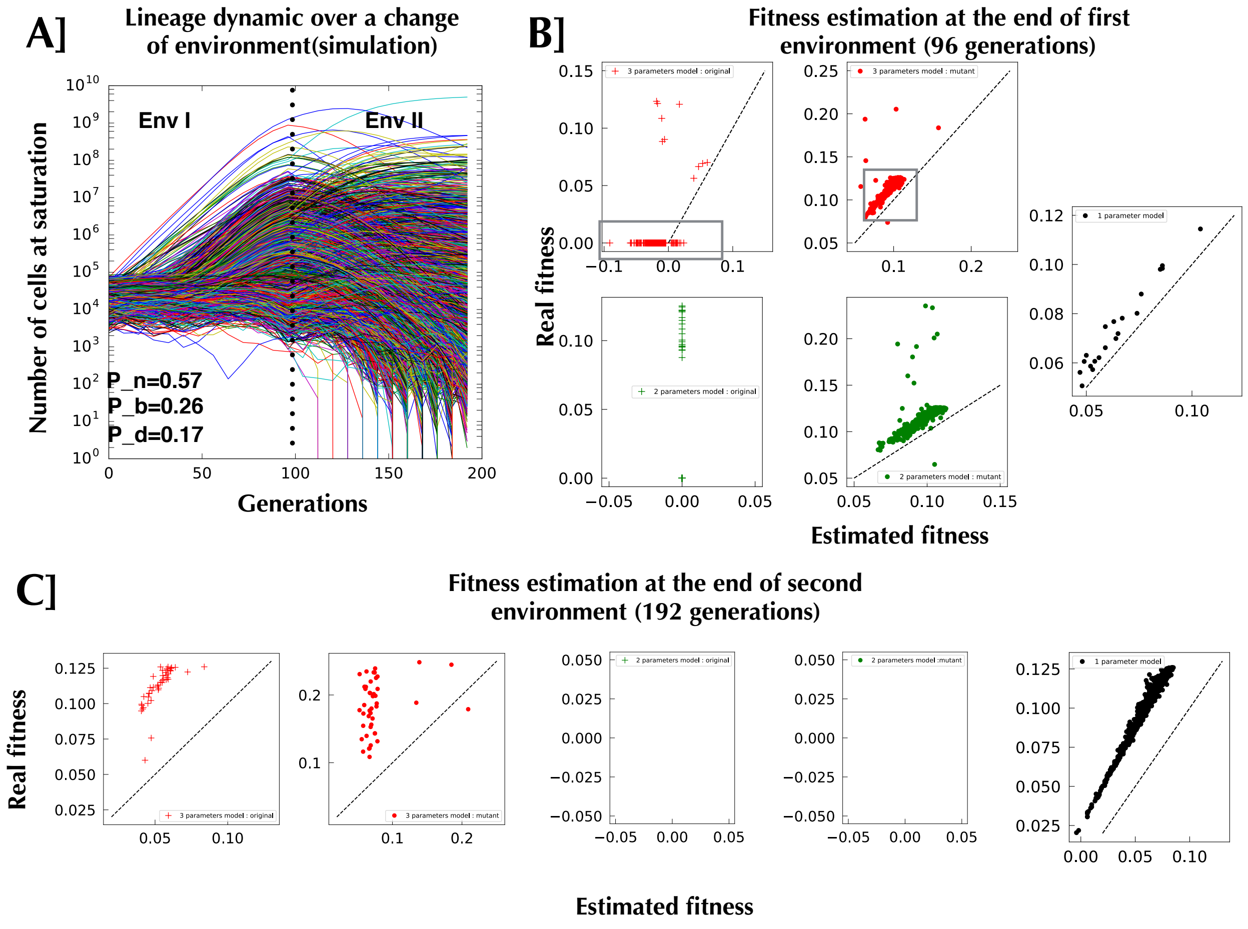


**Figure SI 18 Parameter estimation from simulation with lineage mean fitness criterion and with population mean fitness calculated from thread. All fitness are per generation. A] Sample of lineages from a simulated lineage tracking experiment**. **B] Fitness comparison between real fitness fed to the simulation and Maximum Likelihood estimation of fitness for the first environment.** The estimation includes only the top 1,000 lineages present at the end of each environment. It is to be noted that for the 2 parameters model original: a vast majority of the points are by definition stacked at coordinate (0,0), the others, only representing dozens of poorly estimated points out of 1,000. The discrepancy between estimated fitness and real fitness emphasized by the grey box, can be explained. Those boxes contain the same lineages. In the simulation for those lineages, there is no second mutation on the background of an original mutant: the two mutants coexisted before the start of the first environment because of the probability to mutate during the 16 generations of library preparation. They are not related to each other. The algorithm still predicts the correct fitness (has seen in the second upper panel of B) but was unable to choose the best model. Adding another model where the two mutants exist in the lineage before the environment where we performed the measurement does not solve the problem Fig.SI. 21 and 22. **C] Fitness comparison between real fitness fed to the simulation and ML estimation of fitness for the second environment.** The estimated fitness and the true fitness are well correlated but there is an offset due to the way we calculated the mean fitness. The discrepancy between real fitness and estimated fitness in the second panel comes from the fact that those mutants rose in the last ten generations of the simulation. In other words they should not count: the algorithm should have chosen a one parameter model instead of the three parameters model. Another possibility is that some mutations actually arose but then were outcompeted by the growing population mean fitness. In that case the simulation cannot see it at the final time point, but a 3 parameters model was necessary because at some point this mutant contributed to the mean fitness of the lineage.

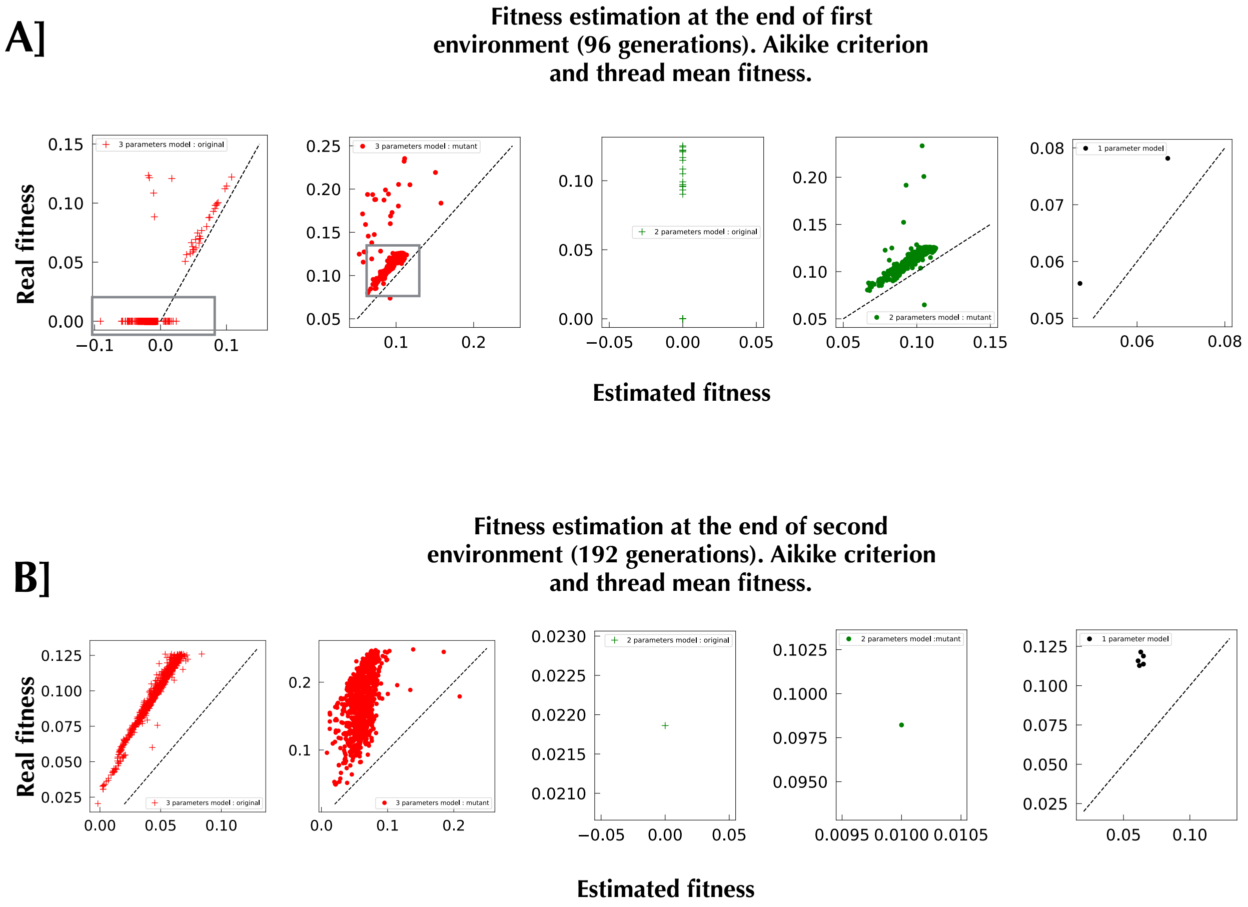


**Figure SI 19 Parameter estimation from simulation with Aikike criterion and with population mean fitness calculated from thread.** As above, but with an AIC for model differentiation.

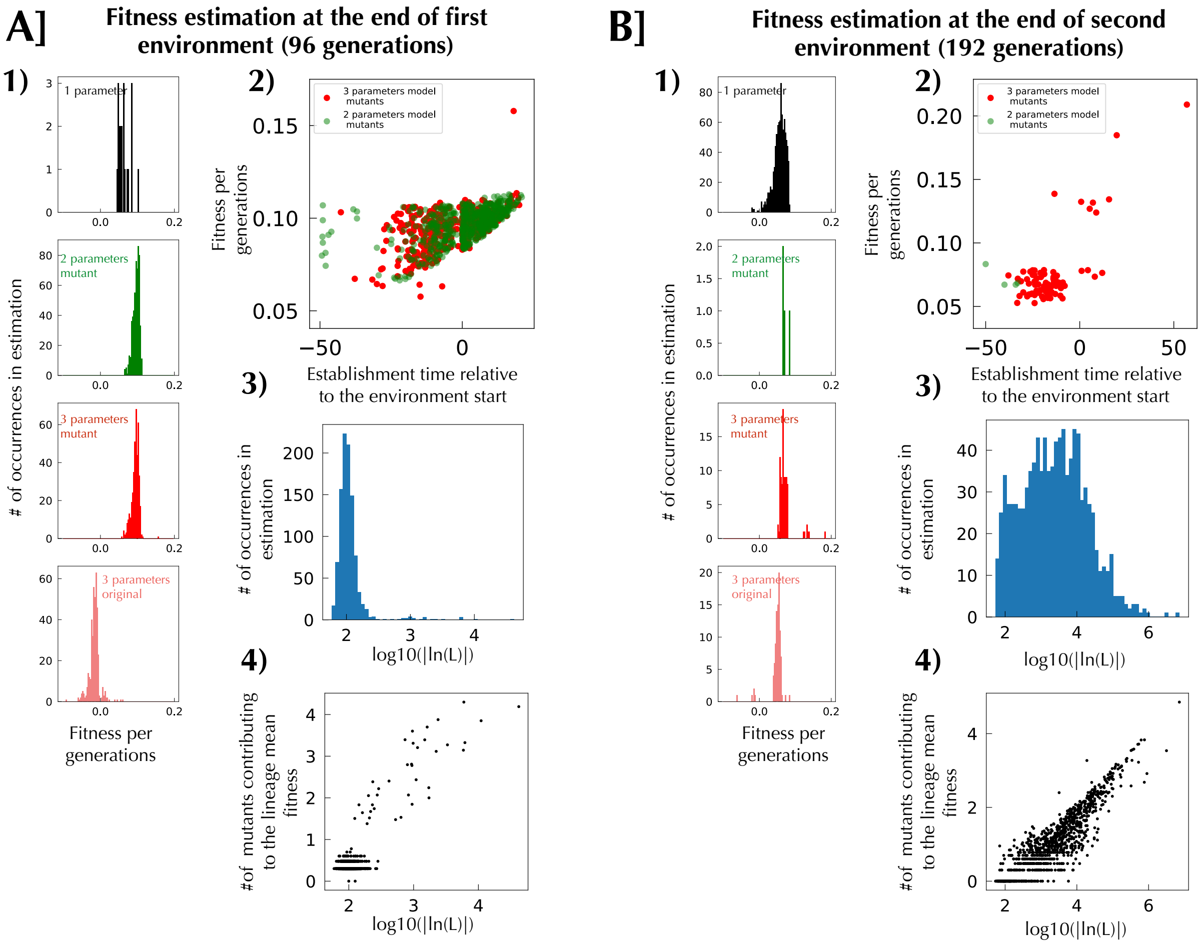


**Figure SI 20 Simulation: full analysis from lineage mean fitness criterion and with population mean fitness calculated from thread. A] Analysis for the first environment.** 1) Distribution of fitness effect according to the different model picked. 2) Space phase for evolution in the first environment. We can see that some of the fitness have an establishment time more negative than that allowed by their fitness. We checked if a model which would represent the trajectories as two existing mutants sharing the lineage at time zero, could alleviate this observation, but saw no improvement (Fig.SI 21 and 22. 3). Distribution of log likelihood for picked models. 4) Relation between the number of mutants that participate to the fitness of a lineage (non-zero at the end of the environment) and the goodness of the fit for those lineages: the worst estimations are well correlated with the hypothesis of the single mutant being broken. **B] Analysis for the second environment**. Same as **A]** but the goodness of fit 3) is now orders of magnitude worse than in panel A] and the trend with the number of mutants is even clearer. The relative poorness of the fit can be explained by our crude estimation of mean fitness, as we can see that goodness of fit comes back to acceptable level when a better estimation of mean fitness is used Fig.SI 24 and 25.

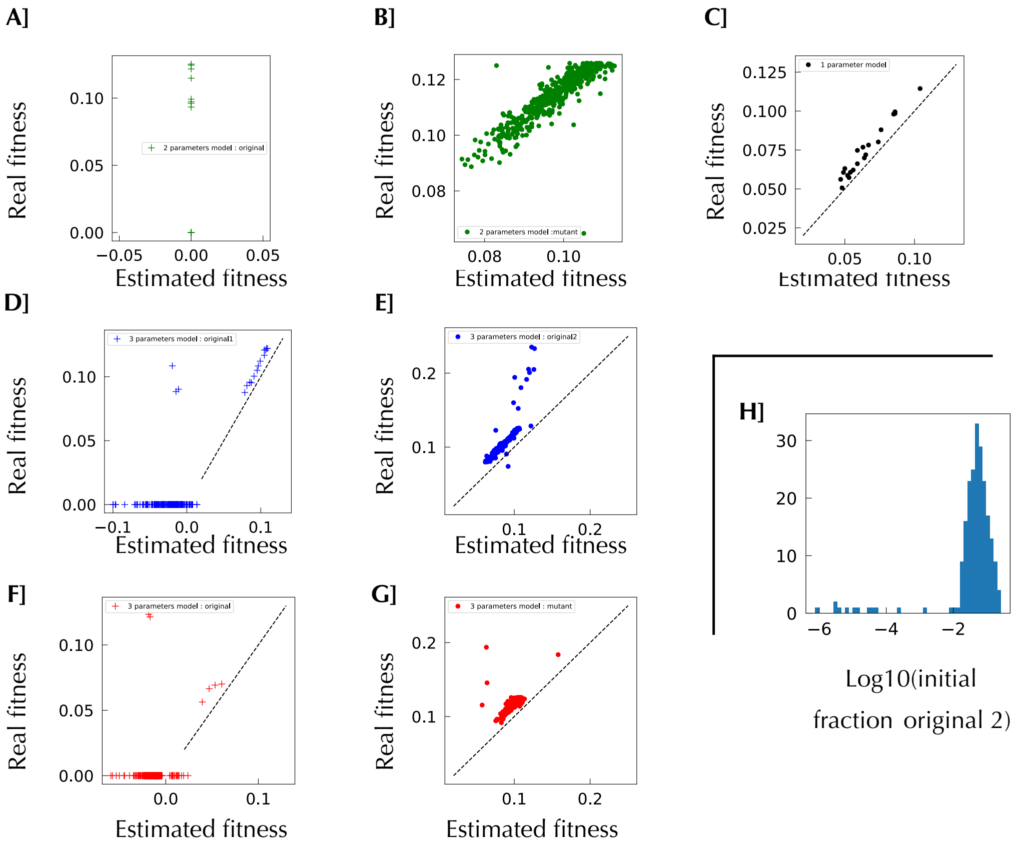


**Figure SI 21 Simulation first environment: Comparison between model for fitness estimation and real fitness from the simulation with additional model: using lineage mean fitness criterion and with population mean fitness calculated from thread. A] 2 parameters model: Original. B] 2 parameters model: Mutant.** As we removed the estimation for which the establishment time was not coherent with the fitness we see a better correlation between estimation and reality. Nonetheless, those removed points have to be estimated with the initial mix model which is a model that assume that two mutants are present in the lineage at the start of the experiment. **C] One parameter model. D] 3 parameters model initial mix of mutants: original 1 (larger fraction).** This model suffers from the same problems as the 3 parameter model with rising mutant: poor estimation of original 1.**E] 3 parameters model initial mix of mutants: original 2 (smaller fraction). F] 3 parameters model rising mutant: original.** Considering the incoherent fitness to establishment time estimation and estimate those lineages with another model, did not solve our problem: bad estimation of original mutant. **G] 3 parameters rising mutant: mutant. H] Distribution of fraction for the original 2 from initial mix of mutants model.** We can see that on average the initial mix mutant model predicts an initial 5 to 10% of the lineage made of the original 2, which is not negligible.

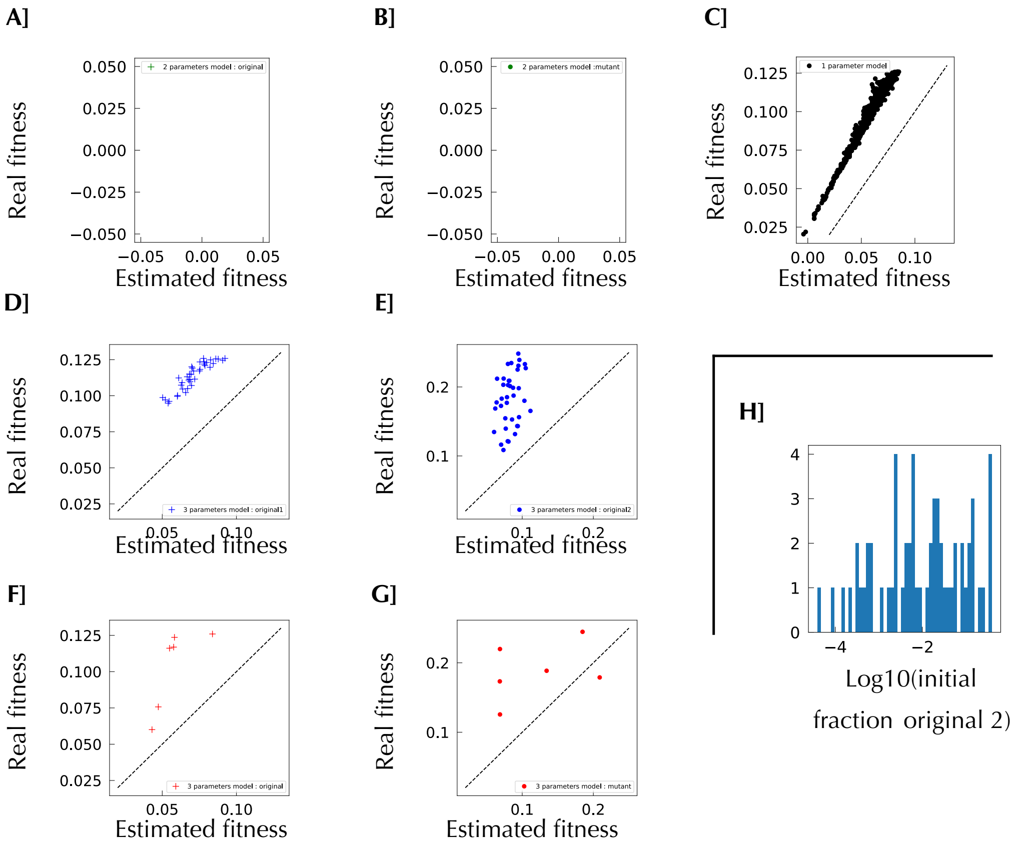


**Figure SI 22 Simulation second environment: Comparison between models for fitness estimation and real fitness from the simulation with additional models: using lineage mean fitness criterion and with population mean fitness calculated from thread. A] 2 parameters model: Original. B] 2 parameters model: Mutant. C] One parameter model. D] 3 parameters model initial mix of mutants : original 1 (larger fraction). E] 3 parameters model initial mix of mutants: original 2 (smaller fraction). F] 3 parameters model rising mutant: original. G] 3 parameters rising mutant: mutant. H] Distribution of fraction for the original 2 from 3 parameters model initial mix of mutants.**

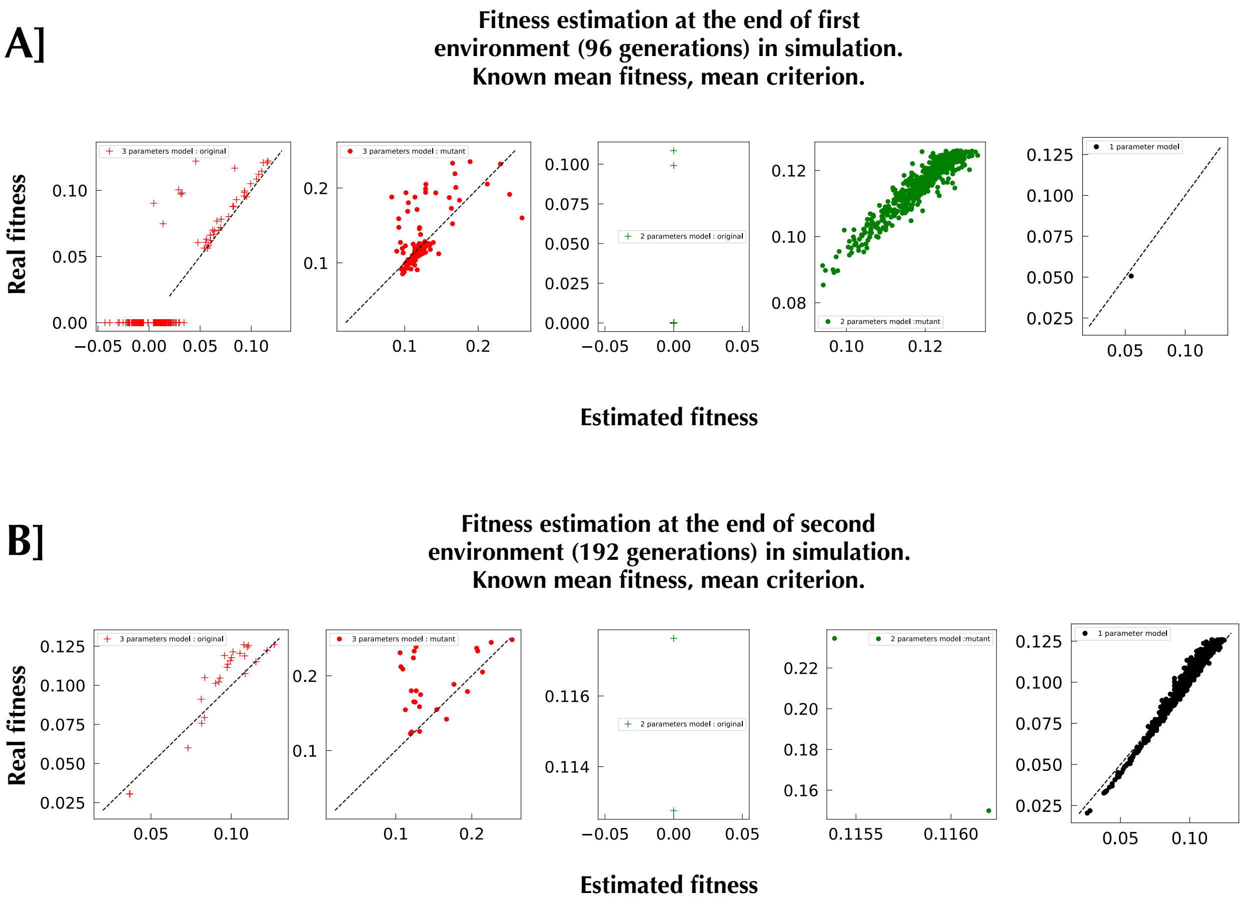


**Figure SI 23 Parameters estimation from simulation with lineage mean fitness criterion and with population mean fitness calculated from known lineage fitness.**

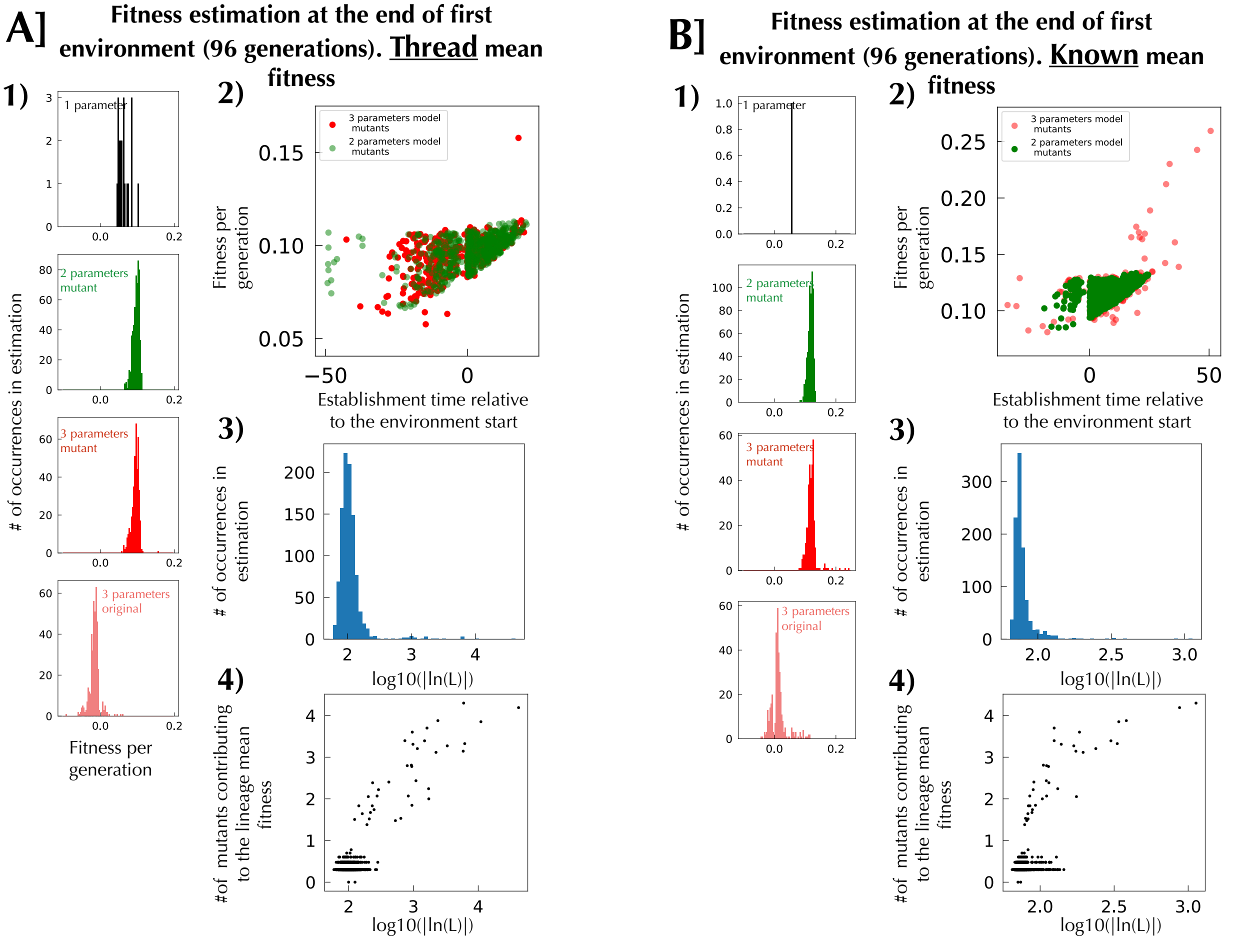


**Figure SI 24 Overall comparison of the fits in environment 1 of simulation, of the first 1,000 largest lineages at the end of environment 1: A] Using for mean population fitness estimation either the thread method or B] Known lineages' fitness.** Using the population mean fitness calculated from known lineage fitness strongly increases the goodness of fit (comparing panels A]3) and B]3)). In terms of distribution of the different parameters there is no striking differences even though the two ways of calculating mean population fitness shows sensible differences.

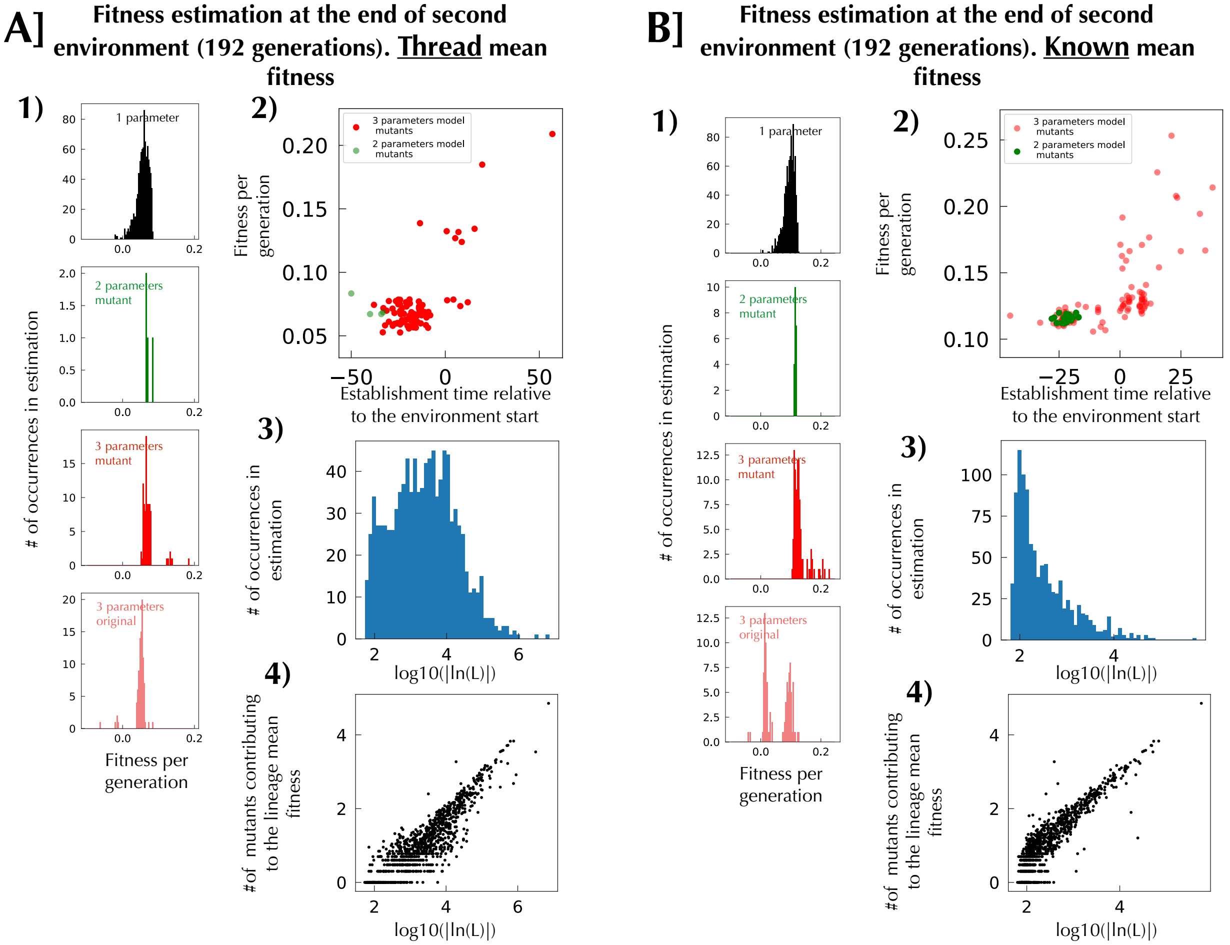


**Figure SI 25 Overall comparison of the fits in environment 2 of simulation, of the first 1,000 biggest lineages at the end of environment 2: A] Using the thread method or B] Known lineages fitness for estimation of the population mean fitness.** Using the mean fitness from known lineage fitness strongly increase the goodness of fit (comparing panels A]3) and B]3)). In terms of distribution of the different parameters there is no striking differences even though the two ways of calculating mean population fitness shows sensible difference Fig.SI 17.

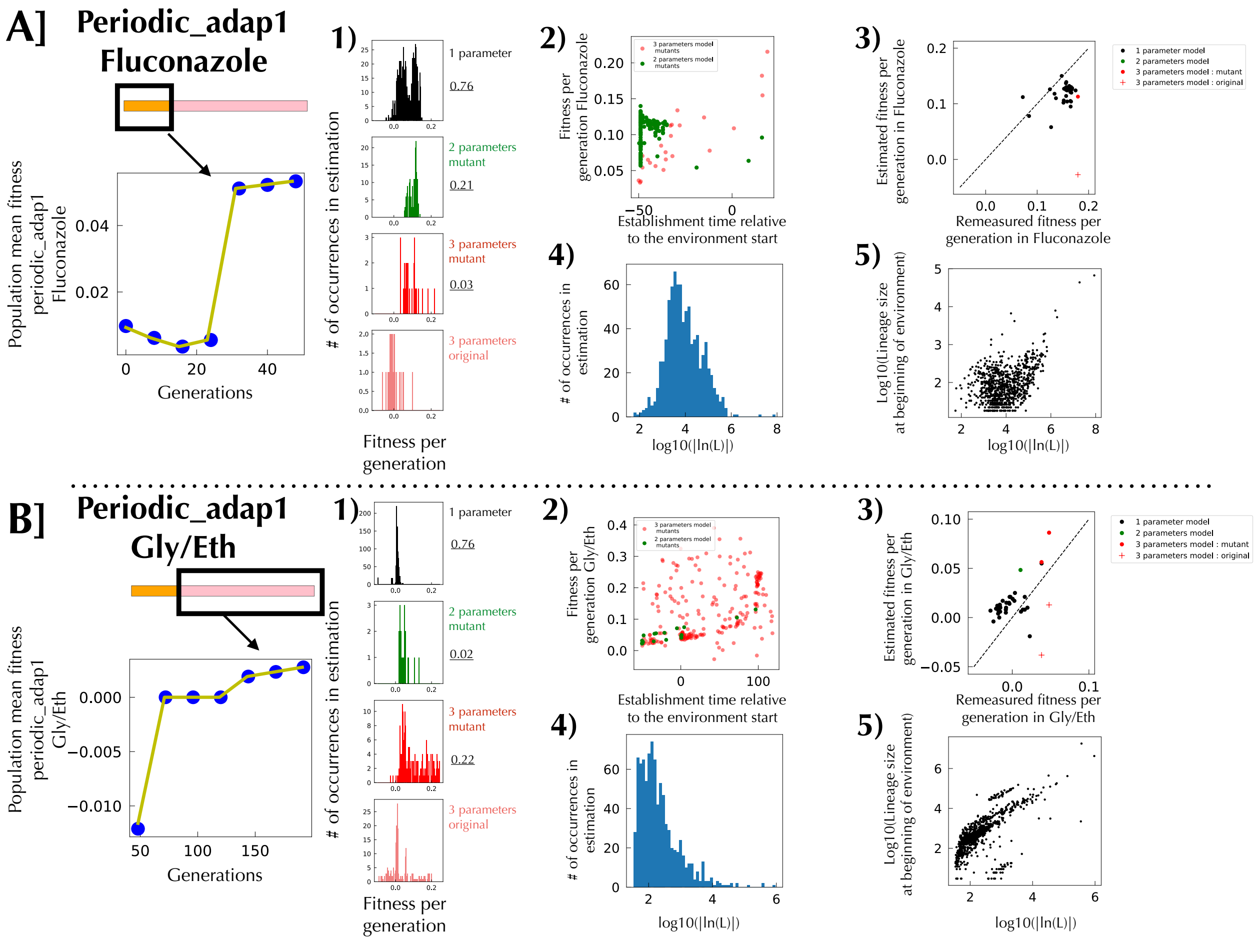


**Figure SI 26 Recapitulation of lineage tracking analysis for periodic_adap1 evolution using a model choice based on lineage fitness criterion and a population mean fitness calculated from thread. A] Analysis of the top 1,000 largest lineages at the end of the first environment.** The most left panel is an estimation of the evolution of the population mean fitness using the exponential decay of lineages behaving similarly (thread). The population mean fitness function has a big jump in the middle, which follows well the type of behavior that we see in the lineage tracking, but which is not a behavior expected for usual population mean fitness. 1) Distribution of fitness effects according to the different model picked. The underlined number on the right of each panel represents the fraction of this particular model chosen by the algorithm. 2) Space phase for evolution in the first environment. 3) Fitness comparison between measured fitness and Maximum Likelihood estimation of fitness for the first environment of periodic_adap1. The estimation follows quite well the measured fitness with of course an offset that comes from our estimation of mean fitness. 4) Distribution of log-likelihood for picked models. The fits are not very good as they are peaked around -10^4 where as good fit usually peaked around -10^2 (see simulations). This is probably coming from the big jump in mean fitness that we cannot explain. 5) Relationship between the size of a lineage at the beginning of an environment (as a proxy for the number of mutants contributing to the mean fitness of the lineage) and the goodness of the fit for those lineages. L is the likelihood of the model. **B] Analysis of the top 1,000 largest lineages at the end of the second environment.** Everything is smoother and makes more sense for the second environment. Still the goodness of fit is quite bad and deserve to be reestimated using population mean fitness estimated from known lineages

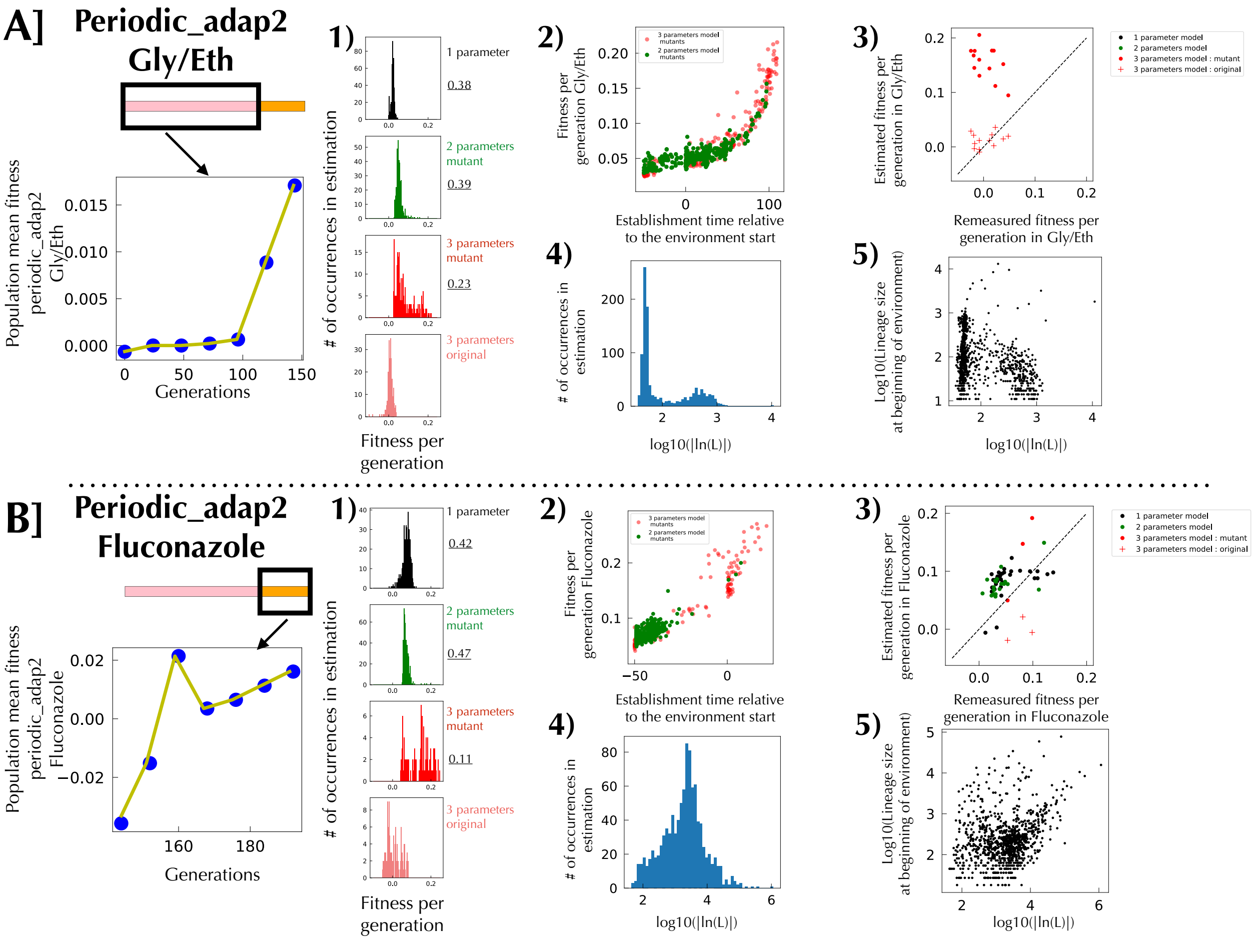


**Figure SI 27 Recapitulation of lineage tracking analysis for periodic adap2 evolution using a model choice based on lineage fitness and a population mean fitness calculated from thread.**

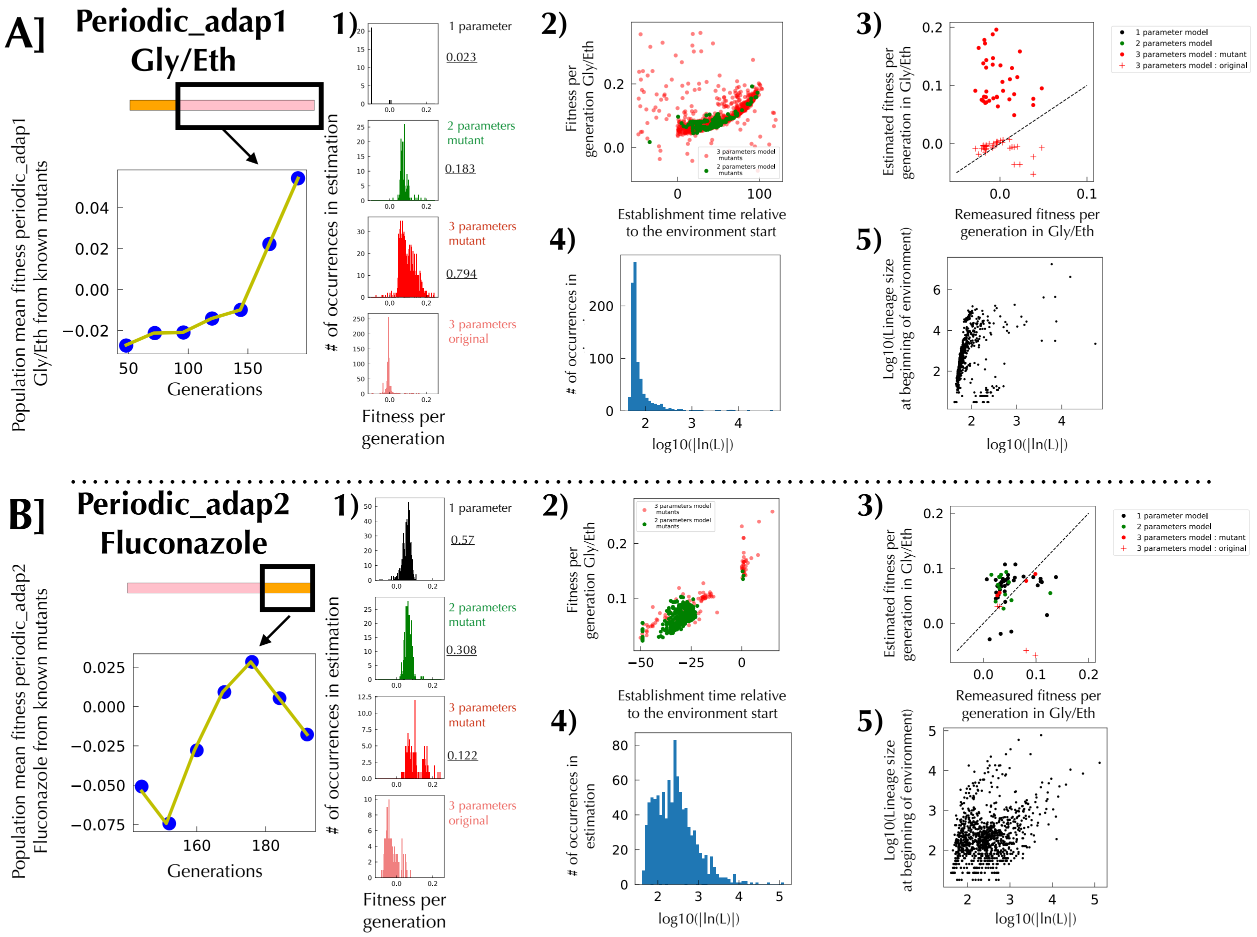


**Figure SI 28 Recapitulation of lineage tracking analysis for second environment of periodic adap1 and 2 evolution using a model choice based on lineage mean fitness and a population mean fitness calculated from known lineages. A]** The goodness of fit distribution 3) is orders of magnitude better than with the thread way to calculate population mean fitness. In addition, most of the 1 parameter models seen before have been moved to a three parameters model. **B]** The goodness of fit distribution 3) is orders of magnitude better than before even though still being quite large. If one add to that the look of the mean population function associated to that Fluconazole environment in all our experiment, it is obvious that we are lacking a full description of that environment.

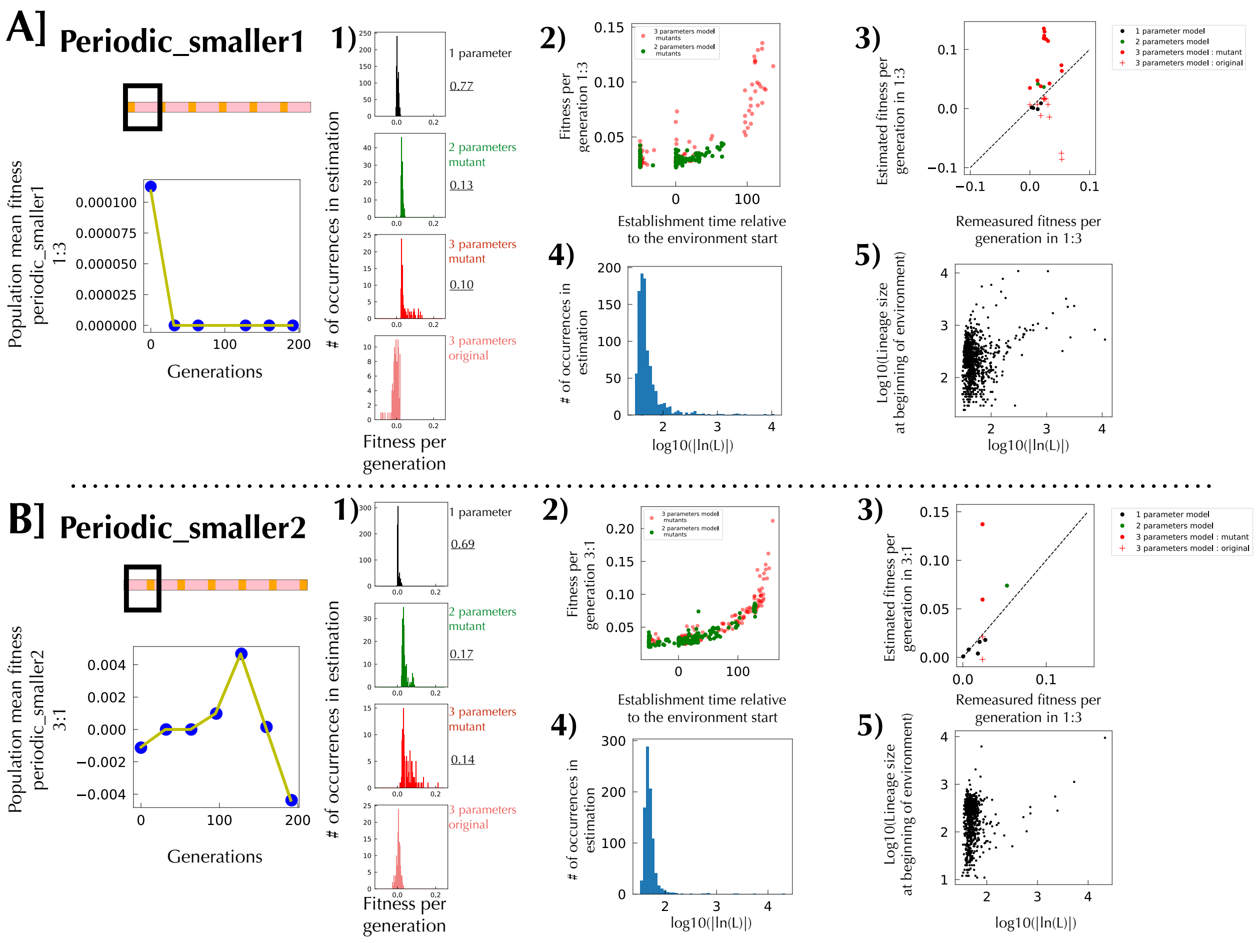


**Figure SI 29 Recapitulation of lineage tracking analysis for periodic adap2 evolution using a model choice based on lineage fitness and a population mean fitness calculated from thread.**

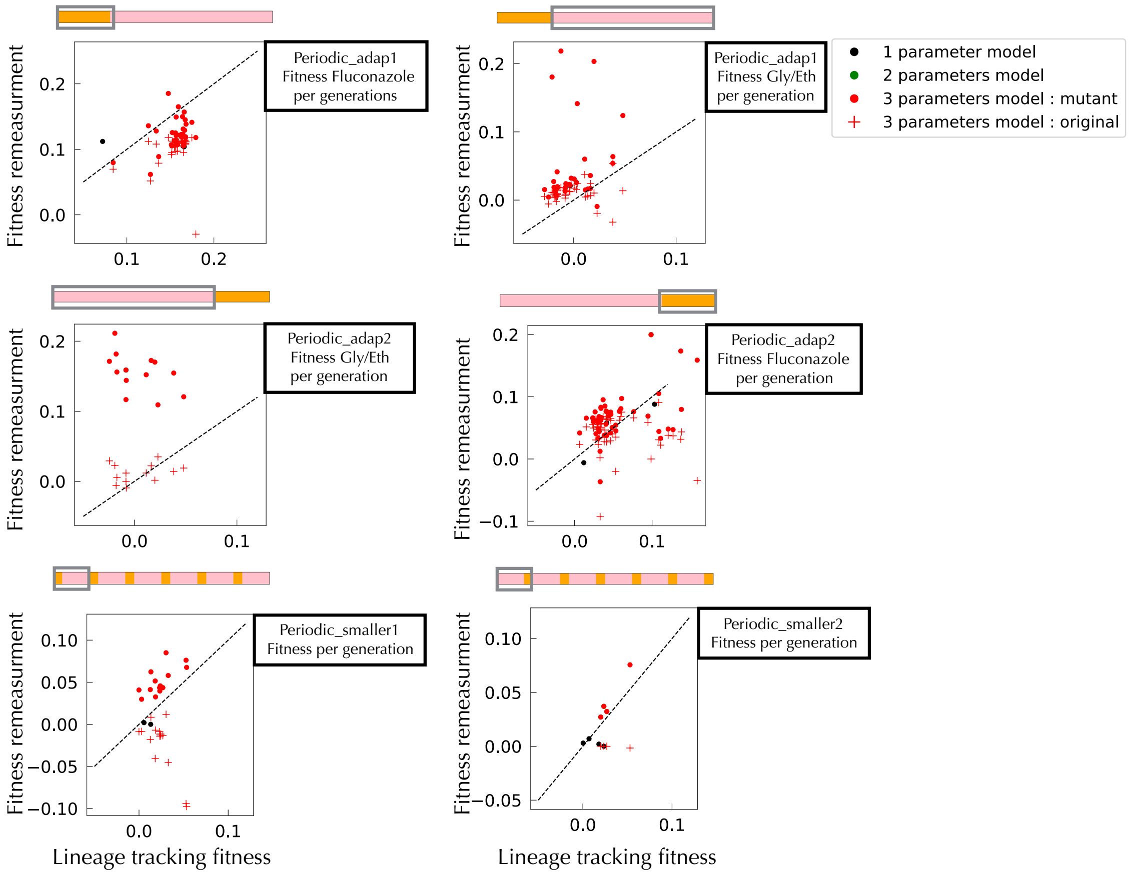


**Figure SI 30 Aikike criterion and thread mean fitness: Comparison of fitness between remeasurement and Maximum Likelihood estimation from lineage tracking.**

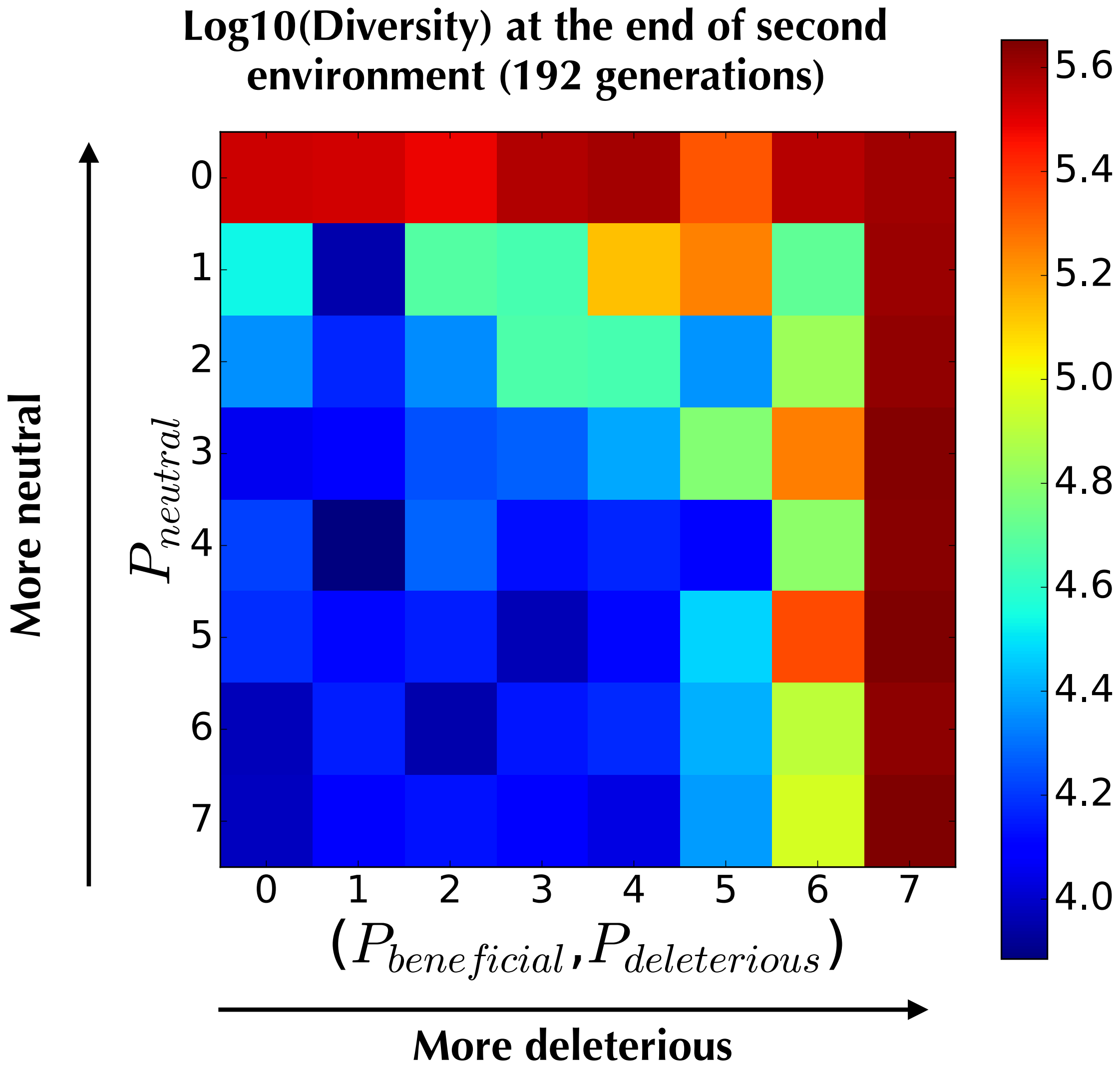


**Figure SI 31: Simulation of barcode diversity loss after 192 generations due to uncorrelated fitness changes between two environments.** Here the two environments have the same uniform DFE ]0,125] ,non synonymous mutation rate mu_env=10^-5 and consecutive spent time of 96 generations. The initial barcode diversity is of 500 000 barcodes. Barcode diversity loss was calculated for simulation were Pn, Pb and Pd were varied. There is no correlated behavior of a mutant between the two environments. We see that unless high bias toward a joint distribution of fitness effect of those two environments that links preferentially a beneficial mutation in the first environment to a deleterious or neutral fitness in the second environment, we end up with a reduction of diversity of 50 fold after 192 generations.

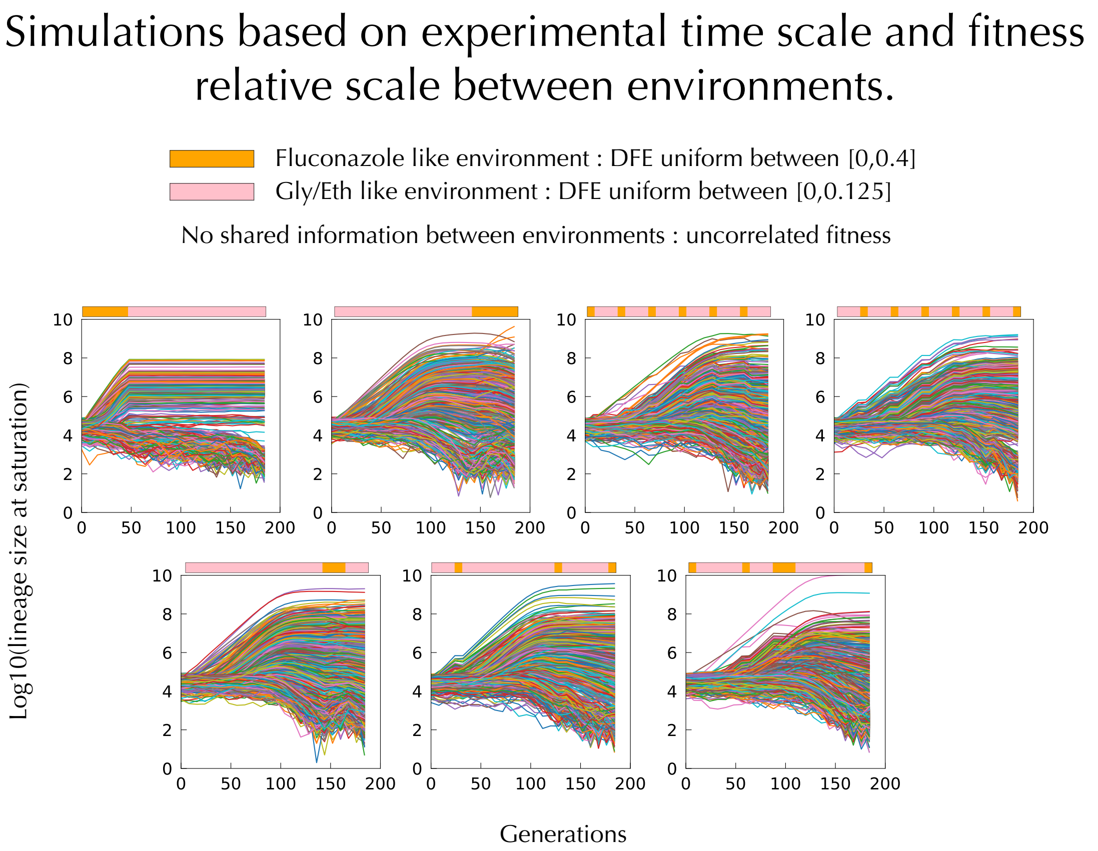


**Figure SI 32: Lineage tracking for a subsample of randomly selected lineages from simulations mimicking our experiments.** In those simulations Pb=Pd=0.25 and Pn=0.5. There is no correlation between fitness in the two environments. Mu_env Gly/Eth=10^-6 and Mu_env Fluconazole=10^-5. The bounds for the uniform DFE respects the 3 fold scale difference and range between fitness in Fluconazole and in Gly/Eh observed experimentaly : respectively ]0,0.125] per generation (]0,1] per cycle) and ]0,0.05] per generation (]0,0.4] per cycle). Those parameters allows to recover the 20% of adaptation after 48 generations in Fluconazole but failed to recover the 15 % of adaptation for Gly/Eth. Those simulations show very different behavior from what we observed, thus showing the importance of the form of the DFE , the joint distribution of fitness effect and target size of mutations, to be predictive of the dynamic of adaptation in changing environment.
